# Harnessing machine translation methods for sequence alignment

**DOI:** 10.1101/2022.07.22.501063

**Authors:** Edo Dotan, Yonatan Belinkov, Oren Avram, Elya Wygoda, Noa Ecker, Michael Alburquerque, Omri Keren, Gil Loewenthal, Tal Pupko

**Author notes:** To whom correspondence should be addressed: Tal Pupko.

## Abstract

The sequence alignment problem is one of the most fundamental problems in bioinformatics and a plethora of methods were devised to tackle it. Here we introduce BetaAlign, a novel methodology for aligning sequences using a natural language processing (NLP) approach. BetaAlign accounts for the possible variability of the evolutionary process among different datasets by using an ensemble of transformers, each trained on millions of samples generated from a different evolutionary model. Our approach leads to outstanding alignment accuracy, often outperforming commonly used methods, such as MAFFT, DIALIGN, ClustalW, T-Coffee, and MUSCLE. Notably, the utilization of deep-learning techniques for the sequence alignment problem brings additional advantages, such as automatic feature extraction that can be leveraged for a variety of downstream analysis tasks.

## Main Text

A multiple sequence alignment (MSA) provides a record of homology at the single position resolution within a set of homologous sequences. In order to infer the MSA from input sequences, one has to consider different evolutionary events, such as, substitutions and indels (i.e., insertions and deletions). MSAs can be computed for DNA, RNA, or amino acid sequences. MSA inference is considered one of the most common problems in biology (*1*). Moreover, MSAs are required input for various widely-used bioinformatics methods such as domain analysis, phylogenetic reconstruction, motif finding, and ancestral sequence reconstruction (*2*). These methods assume the correctness of the MSA, and their performance might degrade when inaccurate MSAs are used as input (*3, 4*).

MSA algorithms typically assume a fixed cost matrix as input, i.e., the penalty for aligning non-identical characters and the reward for aligning identical characters. They also assume a penalty for the introduction of gaps. These costs dictate the score of each possible alignment, and the algorithm aims to output the alignment with the highest score. MSA algorithms often allow users to select among several default scoring schemes and tune parameters that control the MSA computation. However, in practice, these parameters are seldom altered, and only a few default configurations are used. Previous studies demonstrated that fitting the cost matrix configuration to the data increases the MSA inference accuracy (*5, 6*).

In the last decade, deep-learning algorithms have revolutionized various fields (*7*), including computer vision (*8*), natural language processing (NLP) (*9*), sequence correction (*10*), and medical diagnosis (*11, 12*). Neural-network solutions often resulted in a substantial increase in prediction accuracy compared to traditional algorithms. Deep learning has also changed molecular biology and evolutionary research, e.g., by allowing accurate predictions of three-dimensional protein structures using AlphaFold (*13*).

To the best of our knowledge, machine translation is not currently used to align sequences. We propose BetaAlign, a novel deep-learning method that is trained on known alignments, and is able to accurately align novel sets of sequences. The method is based on a recent state-of-the-art deep-learning architecture for machine translation called “transformer” (*14*). BetaAlign was trained on millions of alignments drawn from different evolutionary models. We analyze the accuracy of BetaAlign and show that it is comparable, and in some cases superior to the most popular MSA algorithms, T-Coffee (*15*), ClustalW (*16*), DIALIGN (*17*), MUSCLE (*18*), MAFFT (*19*) and PRANK (*20*).

BetaAlign converts the alignment problem into a translation problem, a well-studied problem within the NLP field (*21*). To frame an alignment task as an NLP translation task, we represent both the unaligned sequences and the resulting alignment as sentences in two different “languages”. We first present the NLP approach for pairwise alignment. We represent the two input sequences as a sentence in a language that we term “concat” (the source language). It is termed “concat” because, in this representation, each pair of input sequences is concatenated, with the pipe character (“|”) representing the border between the sequences. In this language, each character in the resulting string is considered a different word. For example, when we aligned the sequence “AAG” against the sequence “ACGG”, in the “concat” language, this will be represented by the sentence “A A G | A C G G” (Fig. 1). In this sentence, there are eight words. Of note, in the “concat” language there are only five possible unique words. A dictionary of a language is the entire set of words used in that language, and thus in the “concat” language, the dictionary is the set {“A”, “C”, “G”, “T”, “|”}.

**Fig. 1.**
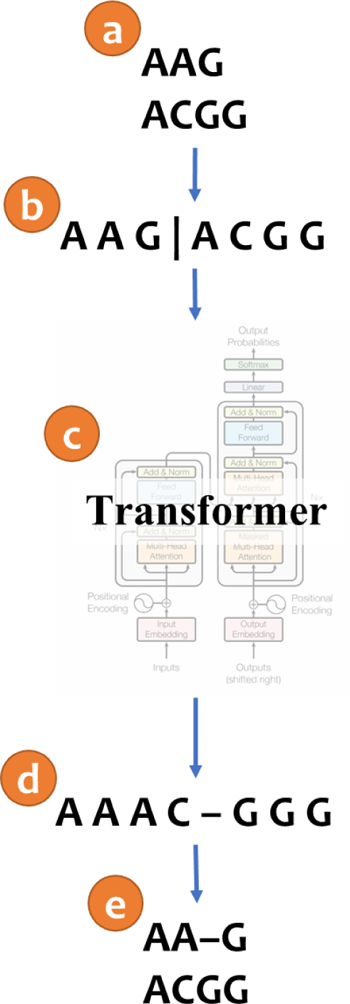
An illustration of aligning sequences with the machine translation method. (a) Consider two input sequences “AAG” and “ACGG”. (b) The result of encoding the unaligned sequences into the source language (“concat” representation). (c) The sentence from the source language is translated to the target language via a transformer model. (d) The translated sentence in the target language (“spaces” representation). (e) The resulting alignment, decoded from the translated sentence, in which “AA-G” is aligned to “ACGG”. The transformer architecture illustration is adapted from (14).

The target language, i.e., the language of the output alignment, is termed here “spaces”. In this language, the dictionary is the set {“A”, “C”, “G”, “T”, “–”}. In this representation, the words of the two different aligned sequences are interleaved. That is, the first two words in the “spaces” output sentence are the first character of the first and the second sequence, respectively. The third and fourth words correspond to the second column of the pairwise alignment, etc. In the above example, assume that in the output alignment “AA-G” is aligned to “ACGG”, then, in the “spaces” language, this will be represented as the sentence “A A A C – G G G” (Fig. 1).

With these representations, the input sequences are a sentence in one language, the output alignment is a sentence in another language, and a perfect alignment algorithm is requested to accurately translate one language to the other. The details regarding the training and execution of the NLP transformers are provided in the Supplementary Text.

We compared the performance of BetaAlign to the state-of-the-art alignment algorithms: ClustalW (*16*), DIALIGN (*17*), MAFFT (*19*), T-Coffee (*15*), PRANK (*20*), and MUSCLE (*18*). For each number of sequences from two to ten, the performance was compared on a simulated test dataset comprising 3,000 nucleotide MSAs (Fig. 2a). BetaAlign had the lowest error rate for any number of aligned sequences (paired t-test; P < 10^−20^). For all alignment methods, the error increases as the number of sequences increases. Notably, ClustalW and DIALIGN were much more affected by the increase of the number of input sequences, compared to other methods. Of note, this analysis was conducted by applying the “concat” source language with the “spaces” target language (see Supplementary Text).

**Fig. 2.**
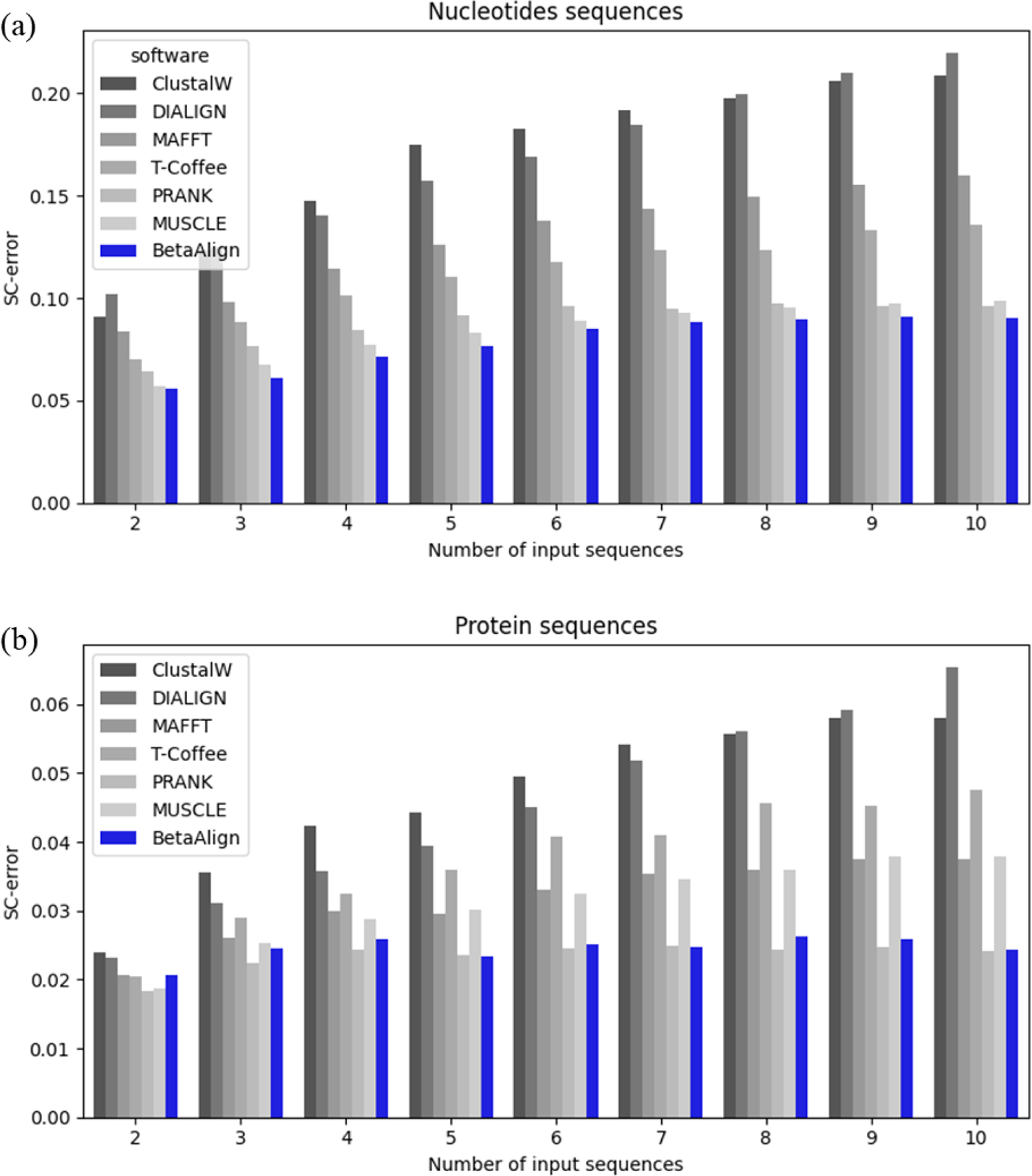
Comparing the accuracy of inferred alignments of ClustalW, DIALIGN, MAFFT, T-Coffee, PRANK, MUSCLE, and BetaAlign. Nucleotide (a) and protein (b) sequences were analyzed on datasets SND1 and SPD1LG, respectively. The x-axis is the number of input sequences, and the y-axis is the SC-error, which quantifies accuracy (see Methods).

We repeated the same analysis for the protein dataset. The error of all methods was low compared to nucleotide sequences, most likely reflecting the increase in alphabet size, which helps identify homologous regions (Fig. 2b). PRANK and BetaAlign outperformed all other aligners. As seen from Fig. 2, there is no single optimal method to align all types of sequences.

In order to further improve BetaAlign’s performance (see Supplementary Text), we have used: (1) transfer learning, which allows optimizing a transformer based on previously trained transformer weights; (2) focus learning, which optimizes the performance on a subspace of the alignment parameters space. To further characterize BetaAlign, we conducted the following analyses: (1) evaluating the effect of training time and size; (2) measuring the performance as a function of the evolutionary dynamics that generated the sequences; (3) evaluating the sensitivity to model misspecification; (4) comparing different encoding representations; (5) comparing different transformer architectures; and (6) analyzing the attention mechanism, which highlights how alignment transformers differ from those used to translate natural languages (see Supplementary Text).

In a broad perspective, the transformer modular architecture allows replacing the decoder with alternative task-specific architectures, for instance, classification or regression. The weights that were learned by the encoder could be used as a starting point for other machine-learning models (as it was already trained on millions of genomic sequences). More concretely, we show that our approach also provides the benefit of automated extraction of features (from raw unaligned sequence data) that can be used for predicting the length of ancestral sequences (see Supplementary Text).

Of note, several hurdles were encountered using the development of BetaAlign, e.g., computational challenges associated with aligning long sequences and the occasional generation of invalid alignments. Both these challenges were addressed by introducing modifications to the NLP algorithms. All these issues are detailed in the Materials and Methods and Supplementary Text.

We have designed and implemented a new technology that aligns sequences with accuracy that competes with state-of-the-art methods. Our deep learning-based approach is fundamentally different from current algorithms, which rely on conventional scoring schemes. Our method allows us to personalize the alignment process for inputs that differ in their evolutionary dynamics. The increased knowledge regarding the differences in mutation processes and selection regimes among different species allows us to easily train BetaAlign on clade-specific datasets, i.e., we are capable of creating different aligners per kingdom and even per species. Similarly, BetaAlign can be trained to capture the evolutionary dynamics of different regions within the genome, e.g., introns versus promotors versus other non-coding regions. Moreover, this method has the potential to capture variations among different types of protein-coding genes, e.g., transmembrane versus cytoplasmatic proteins.

One can argue that BetaAlign heavily depends on the simulator. In other words, if the simulator does not reflect biological realism, the resulting aligner may produce inaccurate MSAs for empirical data. However, the design of BetaAlign allows switching between simulators easily. This allows research groups to focus on mimicking the sequence evolutionary process to create better simulators, which in turn, will improve the alignment accuracy. Since tuning a simulator is an easier task than creating a new alignment method, we anticipate a shift in the scientific community from developing better aligners to developing more realistic simulation schemes. BetaAlign can learn those specific alignment rules without the need to specifically provide them as an input to the program. This feature abolishes the need to provide any assumptions regarding the evolutionary process at the development stage of the aligner. For example, if a research group assumes a translocation or recombination happened between proteins of different species, they can create a simulator reflecting this process and train BetaAlign to align those sequences, instead of developing new methods to consider translocation or recombination (to the best of our knowledge, current aligners do not consider these types of events).

In this pioneering report, we have demonstrated the power of applying NLP tools to biological questions. We believe that this link is essential and will change many aspects of computational biology. Leveraging deep-learning models discovers new research options with promising results. Of course, the NLP methods could be used in many other biological tasks, e.g., one could translate unaligned sequences to generate the ancestor sequence or classify different protein sequences to their corresponding GO annotations. Many downstream tasks could benefit from accurate features which could be extracted by embedding the genomic data.

We have coupled the NLP domain and the multiple sequence alignment problem by using transformers that were originally designed for natural languages. Thus, future improvements in the NLP field are likely to have a direct impact on future alignment methodologies. We expect that in the next few years, transformers that are dedicated to the task of sequence alignment, together with other breakthroughs in machine learning, will lead to alignment algorithms that account for the specific grammar rules of each set of analyzed sequences and will substantially outperform existing aligners.

## Supplementary Text including Material and Methods

### Materials and Methods

#### Previous Approaches for Aligning Sequences and Motivation for the Current Algorithm

The Needleman–Wunsch algorithm was the first to use dynamic programming to efficiently find the best global scoring alignment among two sequences (*22*). MSA inference was shown to be an NP-complete problem (*23*), making the inference task impractical for a large set of sequences. To overcome this hurdle, popular MSA algorithms, such as MAFFT (*19*) and PRANK (*24*), use heuristics to reduce the search space and consequently, the running time.

There is extensive knowledge regarding the variability of the evolutionary process among different datasets and lineages. For example, amino-acid replacement matrices vary between proteins encoded in the nuclear genome, the mitochondria, and plastids (*25*). Indel dynamics also highly vary between datasets and among different phylogenetic groups (*26*). Site-specific evolutionary rates also vary along the analyzed sequence, for example, amino-acid sites that are exposed to the solvent tend to have higher evolutionary rates compared to buried sites (*27*). Alignment algorithms using default configurations implicitly assume that the evolutionary dynamics do not vary among different datasets and within a single dataset. The general inability of MSA inference algorithms to automatically tune their scoring scheme to the specific dataset being analyzed is a shortcoming of present alignment programs. This “one matrix fits all biological datasets” assumption employed by current methodologies raises fundamental questions about the correctness of alignments produced by such methods.

Alignment algorithms are often benchmarked against empirical alignment regions, which are thought to be reliable. However, such regions within alignments do not reflect the universe of alignment problems (*28*). Of note, these regions were often manually computed, so there is no guarantee that they represent a reliable “gold standard” (*29*). When testing alignment programs with simulated complex alignments, the results differ from the benchmarks results (*30*).

One of the key concepts in learning algorithms, in general, and in deep-learning algorithms in particular, is the ability to learn from previously annotated data, i.e., to generalize from previous observations to unseen cases. For the task of alignment inference, a deep-learning algorithm should learn from “true” alignments (e.g., simulated sequences for which the correct alignment is known) and apply the obtained knowledge to align novel sequences. In this work, we demonstrate the use of machine translation to align biological sequences. Our main assumption is that by utilizing NLP models we can learn to capture the evolutionary dynamics of biological sequences.

#### New Approaches

##### An NLP-Based Translation Model

Our hypothesis is that cutting-edge NLP tools can learn from known examples, and a trained model can infer accurate alignments for unseen data. A plethora of neural network architectures is available in the NLP domain for translation, including recurrent neural networks (*31*) and transformers (*14*). As these architectures are sequence to sequence, the input to these models is a sentence, and the output is a predicted translation. Our approach relies on optimizing the model based on training data, i.e., the learning phase of the algorithm. In our case, a massive amount of training data, i.e., true alignments, is obtained using simulations. We simulate the training alignments using SpartaABC (*26*). SpartaABC allows simulating datasets along a phylogenetic tree, with various indel-length distributions, and with different indel-to-substitution rate ratios. It allows finding indel model parameters that best describe a specific empirical dataset. It also assumes that the evolutionary dynamics of insertions and deletions are characterized by a different set of parameters, i.e., a rich-indel model (*26*).

Formally, let *x* denote the input sequences and *y* denote the correct MSA. Each training data point is a pair (*x*, *y*). During the training phase, the NLP algorithm learns a function *f*_θ_ that maps *x* to *y*, i.e., a transformer. This function should minimize the discrepancy between *y* and *f*_θ_(*x*). The function *f*_θ_ depends on a set of tunable parameters, θ, which defines the architecture and the internal representation of the transformer. Given an unseen test sequence, which was not used for training, the predicted alignment is obtained by applying *f*_θ_ to this input.

Note that the values of the pairs (*x*, *y*) depend on an evolutionary model ψ that includes the topology and branch lengths of the phylogenetic tree, as well as the substitution and indel model parameters. Changing ψ would alter the training data distribution, which in turn, should lead to a different *f*_θ_. It is possible to train the model on data from a specific ψ, or train a model in which ψ is sampled from a prior distribution over its model parameters.

If we apply the “concat” encoding with the “pairs” decoding (see Comparing Different Encoding and Transformers Architectures), the source dictionary size is the set of five characters and the size of the target dictionary is the set of 24 characters discussed above. In NLP translation, each word in each dictionary has to be embedded in a higher dimensional space. Thus, two different embeddings are needed (for the source and target dictionaries). In contrast, we can consider a joint dictionary for applying the “concat” or “crisscross” (see Comparing Different Encoding and Transformers Architectures) encodings with the “spaces” decoding. The joint dictionary of “concat” and “spaces” has six words: {“A”, “C”, “G”, “T”, “–”, “|”}. We note that the word “–” is never found in the input, while the word “|” is never found in the output. Nevertheless, we assume a joint dictionary for both the input and the output. The joint dictionary of “crisscross” “spaces” has five words: {“A”, “C”, “G”, “T”, “–”}. Thus, the same embedding function is used for both the source language, i.e., the unaligned sequences, and the target language, the aligned sequences. One of the advantages of the joint dictionaries explained above over the separate dictionaries of “concat” with “pairs”, is that the dictionaries have a fixed size regardless of how many sequences are included in the alignment (in the “pairs” representation, we should use triplets for alignments with three sequences, quadruplets for alignments with four sequences, etc.) Thus, we only use the “spaces” language when we generalize from pairwise to multiple alignment.

##### Coverage

When the trained BetaAlign is applied to test data, the output is a sequence of words. It is possible for this sequence of words to not reflect a biologically meaningful alignment. For example, while an alignment algorithm, in general, should only introduce new gap characters, here, our algorithm sometimes wrongly mutates the characters of the input sequence or outputs an alignment in which the sequences are not of the same length (although, by alignment definition, they should be). We call such an output an “invalid alignment”. Such cases are illustrated in Fig. S1. A detailed explanation about this metric is at the “Calculating Coverage” section below.

It is possible to train several transformers on the same training dataset. These transformers can have different configurations, e.g., as a result of using different optional tunable parameters. One such tunable parameter is, for example, the learning rate (see below). We train a set of transformers on the same training data. It is often the case that one transformer outputs an invalid alignment, while others output a valid alignment. We use only the valid alignments to determine the final output. The performance of BetaAlign is always computed on the output of the original transformer. We term the percentage of cases for which we can obtain valid alignments as “coverage”. The more transformers we use, the higher the “coverage”, and thus the higher our confidence in the final output. Of note, using alternative transformers also provides alternative alignments, which can be used, for example, to estimate alignment uncertainty (*32, 33*).

##### The Transformer Architecture

As transformers are becoming highly popular in the field of NLP and recently also in other fields such as computer vision (*34*), we used them as part of our deep-learning algorithm. A transformer is a deep-learning model designed to handle sequential data. It is based on an attention mechanism, which means that the algorithm assigns more weight to the parts of the input that contribute more to its predictions (*14*). A transformer is composed of two parts: encoder and decoder. The encoder embeds each input sequence into a high-dimensional vector representation. Next, the decoder receives that representation and the last generated word and predicts the next word. We examined several different transformer configurations, mainly exploring three parameters. First, the maximum number of tokens (words) in a batch, i.e., max tokens, which determines the number of learning elements processed before *f*_θ_ is updated. Second, the learning rate, which determines the strength of updating *f*_θ_ in each update cycle, i.e., one forward and backward pass. A high learning rate value may converge faster, but may reach local optima, while low values of the learning rate are more stable but may take many update cycles to reach convergence to optimal parameters. Third, the warmup updates parameter dictates how the learning rate varies during the process of optimizing the transformer. We train the transformer by optimizing first with a low learning rate, and as the number of optimization cycles increases, the learning rate increases as well. In fact, the learning rate increases linearly, until the maximal learning rate (the second parameter) is reached, and then it starts to decrease with a configurable scheduler. The third parameter, in essence, dictates the slope of this linear function. We implemented the transformers using the Fairseq library (*35*) (for the training parameters see Training the Transformers section).

When optimizing the transformer, we use a technique called transfer learning (*36*). In this approach, a model, e.g., a transformer, which was trained for task 1 is the starting model for a related task 2. That is, the initial values of θ for the new learning (task 2) are the optimal ones achieved from task 1. In our problem, we trained one transformer for pairwise alignment, and use its parameters (rather than using random ones) as starting points for training a transformer for aligning three sequences, and so on. We also use transfer learning in order to train a transformer on subregions of the parameters space ψ. For example, we can train a general pairwise alignment transformer, in which the gap distribution follows a Zipfian distribution which is characterized by the power law parameter, alpha, where alpha is sampled uniformly in the interval [1,2). This transformer can be used as the basis for optimizing a transformer in which the alpha parameter is sampled uniformly from the range [1,1.1). It can also be used to optimize a transformer in which the range of alpha is in the interval [1.1,1.2), etc. In essence, this allows training of several transformers, specialized for subregions of the parameter space, i.e., focused learning.

### Generating True MSAs Using SpartaABC

We used SpartaABC (*26*) to generate millions of true alignments. SpartaABC requires the following input: (1) a rooted phylogenetic tree, which includes a topology and branch lengths; (2) a substitution model (amino acids or nucleotides); (3) root sequence length; (4) the indel model parameters, which include: insertion rate (R_I), deletion rate (R_D), a parameter for the insertion Zipfian distribution (A_I), and a parameter for the deletion Zipfian distribution (A_D). MSAs were simulated along random phylogenetic tree topologies generated using the program ETE version 3.0 (*37*) with default parameters.

We describe the data created for Fig. 2 in the main text. We generated 395,000 and 3,000 protein MSAs i.e., SPD1, with ten sequences that were used as training and testing data, respectively. We generated the same number of DNA MSAs i.e., SND1. Each such MSA was simulated along a different random phylogenetic tree. The random topologies were generated using the program ETE version 3.0 (*37*) with default parameters. For each tree, branch lengths were drawn from a uniform distribution in the range (0.5, 1.0). Next, the sequences were generated using SpartaABC with the following parameters: *R_I, R_D* ∈ (0.0, 0.05), *A_I, A_D* ∈ (1.01, 2.0). The alignment lengths as well as the sequence lengths of the tree leaves vary within and among datasets as they depend on the indel dynamics and the root length. The root length was sampled uniformly in the range [32, 44]. Unless stated otherwise, all protein datasets were generated with the WAG+G model, and all DNA datasets were generated with the GTR+G model, with the following parameters: (1) frequencies for the different nucleotides (0.37, 0.166, 0.307, 0.158), in the order “T”, “C”, “A” and “G”; (2) with the substitutions rate (0.444, 0.0843, 0.116, 0.107, 0.00027), in the order “a”, “b”, “c”, “d”, and “e” for the following substitution matrix:

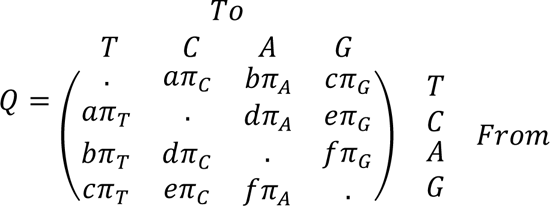

From these ten-taxa trees, we generated trees with fewer sequences. For each alignment, we sampled randomly different number of input sequences to create datasets of 2, 3, 4…10 input sequences. The true alignments that correspond to these trees with fewer taxa were generated by removing the sequences that were not selected from the 10-taxa MSA. It is possible that the resulting MSAs have columns that have only gap characters. These columns were removed to generate valid true MSAs. Of note, before running alignment programs, the resulting sequences in each MSA were unaligned by removing all gap characters from the true MSAs.

Next, we generated a similar amino-acid datasets with a different substitution matrix LG (*38*), which we noted as SPD1LG. Of note, the only differences between SPD1 and SPD1LG is the substitution matrix. The results in Fig. 2 in the main text are on the SPD1LG dataset.

### Generating Long MSAs

For the dataset used in Fig. S6, we generated 395,000 and 3,000 protein MSAs with five sequences, which were used as training and testing data, respectively. We generated the same number of DNA MSAs. As we used segments in those datasets, we divided the alignments into smaller pieces (explained below) with a fixed size of 110. Hence, for the protein dataset the number of sentences in the training data are 2,857,364. For the DNA dataset, the number of sentences in the training data are 2,424,287. The amount of testing segments is not constant as it depends on the transformer results of the previous segment. Of note, when using segments, the number of resulting sentences is higher than the number of alignments (depending on the alignment length and the segment size). In contrast, when aligning shorter sequences, each alignment is a sentence. We used the same SpartaABC parameters as were used for generating the dataset for Fig. 2 (explained above), except for a root length sampled uniformly from the interval [480, 1,120] and a branch length drawn from a uniform distribution in the range (0.1, 0.15). For detailed information about processing the segments dataset, see the Splitting the Unaligned Sequences to Segments section.

### Datasets

Various datasets were generated to train the transformers and to test the performance of BetaAlign. The description of how each data were generated and the parameters used to train each transformer are provided as Supplementary Table S3. The MSAs, trees, and Python scripts used to generate these data are available on GitHub: https://github.com/idotan286/SimulateAlignments.

### Preprocessing the Input for the Machine Translation Task

To train the NLP models, we first represent the input unaligned sequences in one of the two source languages explained above (“concat” or “crisscross”). The true MSAs were also represented in the target languages (“pairs” or “spaces”). The actual training of the NLP transformers as well as the pre-processing step were done using the Fairseq library (*35*), which is implemented in Python. Six text files are used as input for the pre-processing step before actually running the transformer: unaligned sequences and true MSAs, for training, validation, and testing data. Of note, validation data are used to compare between different transformers, which differ in their training parameters such as max token and learning rate. We optimized these training parameters on the first few datasets and then used these optimized parameters for all other runs. Thus, when using the optimized parameters, no validation data were used. The flags used in the preprocessing stage are explained below.

### Preprocessing Inputs and Outputs

In addition, for running the preprocessing steps, one has to declare names for the input and output languages. Moreover, one can explicitly state if the same dictionary is used for both the source and target languages, i.e., whether a joint dictionary should be assumed. In our case, when we apply the combinations of “concat” and “spaces” or “crisscross” and “spaces”, the dictionary is joint, while for the combination “concat” and “pairs”, the dictionary is not joint and hence two different dictionaries are internally generated by the library (one for the source language and the other for the target language). The output of this preprocessing step is binary files that are later used as input to the transformer. This preprocess step allows running different configurations of the transformers (or potentially, other machine-learning models) on the same input files, without the need to perform the same computations on the input files over and over again.

### Training the Transformers

We first assessed various transformer configurations, which differ in their training parameters: max tokens, learning rate and warmup updates. Based on their performance on the pairwise nucleotide and protein alignment datasets (Supplementary Table S3), we selected two optimal transformer configurations, which we call “original” and “alternative”. The learning rate and warmup values for both transformers are 5E-5 and 3,000, respectively. The max token parameter values are 4,096 and 2,048 for the original and alternative transformers, respectively. We used label-smoothed cross entropy (*39*) to compute the loss of the model with a dropout (*40*) rate of 0.3. We used “Adam” (*41*) as the optimizer of the model with 0.9 forgetting factors for gradients and 0.98 for the and second moments of gradients.

All model evaluations are executed on GPU machines, Tesla V100-SXM2-32GB. The Weights and Biases platform (https://wandb.ai) was used to follow the progress of the training steps and helped decide when to stop the learning processes.

### Evaluating Accuracy of Inferred Alignments

Alignment accuracy is often quantified by the column score (CS), which counts the number of alignment columns in the inferred alignment that are identical to the “true” alignment (*42*). We note that we request both the coordinates and the characters of each column to match. For example, consider the case that in a true pairwise alignment, position 17 of the first sequences is an “A”, which is matched to an “A” in position 19 of the second sequences. If in the inferred alignment, position 17 of the first sequence matches position 21 in the second sequence, and they are both “A”, this is not considered a match as the coordinates are not the same. To normalize this score to the range of [0,1], we divided the CS by the total number of columns in the true MSA. The resulting score is termed here SC-score (for the score of columns). The SC-error is defined as 1.0 – SC-score, and thus reflects the level of disagreement between the inferred and true MSAs.

### Transfer Learning Implementation

In our work, transfer learning was repeatedly used for training the transformers. The first protein transformer was trained on a simple dataset of pairwise amino acid sequences (we denote this dataset “PD1”, for protein dataset 1). Its weights were randomly sampled with default values of the Fairseq library. The resulting trained transformer is termed “PT1”, for protein transformer 1. PT1 was next trained on PD2, resulting in PT2, etc. A similar process was used to train the nucleotide-based transformers (NT1, NT2, etc.) on nucleotide (DNA) datasets (ND1, ND2, etc.) Of note, transfer learning was used in this study only when the previous and the currently processed data are encoded using the same representations, i.e., they share the same dictionaries.

### Splitting the Unaligned Sequences to Segments

The transformers we applied for aligning sequences are limited to sentences of up to 1,024 tokens i.e., words, which usually suffices for natural languages. When aligning biological sequences, the sentence that represents the unaligned sequences is often much longer than this threshold, and the transformers fail to translate it due to memory and run-time limitations (both these factors are proportional to the square of the sequence length). We will exemplify our approach for aligning long sequences on an example comprising five sequences, each of which of length of at least 2,000 characters. In our “segmentation” methodology, we first define a parameter that we call segment size, which depends on the number of sequences. In our example of five sequences, we selected a segment size of 110 characters. Thus, when we start the alignment process, we first translate 554 words (110 characters from each sequence and four “|” characters, when using the “concat” representation).

When aligning segments, we use dedicated transformers for this task that we call “segmented transformers”. The segmented transformers share the same architecture with the standard transformers, but they are trained on different data. In addition, while standard transformers were designed to always align the entire input sentence, segmented transformers align only prefixes of the unaligned sequences. Specifically, the segmented transformers were designed so that the last column in the resulting aligned sequences does not include gaps.

To generate the training data for the segmented transformers, we simulated long alignments (see section Datasets). This provides us with long unaligned sequences, which we call *X*, and “true” MSA, which we call *Y*. The training data for the segmented transformer are pairs of (*x*, *y*), in which *x* represents a segment of the unaligned sequence taken from *X*, and *y* represents a subregion of *Y* that corresponds to *x*. Both *x* and *y* are short enough to allow training a transformer.

To divide *X* and *Y* to multiple short (*x*, *y*) pairs, we repeated the following process. We first extract the first 110 characters from each of the five sequences, which will be the unaligned segment, i.e., the *x*. In the true alignment, we find the column with the highest index so that the number of non-gap characters of each sequence to the left of this column is smaller or equal to the segment size. Once this column is found, we inspect if it has any indels. If so, we inspect the previous column. We repeat this process until we find a column without indels. The obtained (first) column without indel characters marks the “true” alignment for this segment, i.e., the *y*. Hence, the source sentence is the first 110 characters of each unaligned sequence (*x*) and the target is the “true” alignment (*y*) from the first column until the indel-free column that was found. From both *X* and *Y*, we remove all the characters that appear in *y* (the characters that were successfully aligned). This process is repeated with the new *X* and *Y* until they are empty, i.e., they were divided to multiple (*x*, *y*) pairs. Of note, when analyzing the last segments, it is possible that fewer than 554 words are translated.

For explanatory purposes, we assume a segment size of three and five input sequences (Fig. S2). Simply diving *X* and *Y* to pairs of arbitrary length introduces various problems as illustrated in Problems Arise When Simply Splitting MSAs section below.

The above procedure described the generation of training data, for which the true alignment is known. We now turn to describe how to run a trained model. Here, we first translate the first segment. However, in order to decide which characters to remove and which to retain, we cannot rely on the identification of an indel-free column in the true alignment as we did for the training data (since we do not know the true alignment). Thus, we analyze the resulting alignment, and mark which characters of the input segments were translated from each sequence. These characters are removed, and the process is repeated. This iteration stops when all characters from the unaligned sequences were translated. Of note, different trained transformers may divide the segments differently. After aligning all of the segments, we reconstruct the resulting alignment by concatenating the resulting alignment of each segment one after another. Coverage and accuracy are measured similarly (as explained for the shorter alignments) after the concatenation.

### Problems Arise When Simply Splitting MSAs

The logic for searching for an indel-free column is illustrated in Fig. S3, i.e., an example to show the problems with not indel-free last column. For explanatory purposes, we assume a segment size of three. The algorithm starts by translating the first three characters from each of the five sequences (Fig. S3b). The indel in the last column of the first segment (Fig. S3c) introduces new complexity, as this indel has a perfect match in the next segment (Fig. S3e). In our approach, this is solved by training segmented transformers. If we choose to use indels in the last column, we move the complexity of the segment approach into the concatenation of the segments afterward. One will need to make sure the indels at the resulted alignment make sense. Of note, this approach makes each of the indels appear in only one segment (but several indels may be in the same segment). Similar cases could happen when the indel splits between two segments.

### Calculating Coverage

Each alignment was passed through both the original and alternative transformers. A valid alignment is defined by the following rules: (1) All alignment rows are of the same length; (2) When removing all gaps from each row of the alignment, the result equals to the unaligned input sequences. For example, for the unaligned inputs: “AAG” and “ACGG”, the inferred alignment, “A-AG” over “ACGG” is valid, whereas for the same input the alignment “A-AG” over “ACGC” is invalid (see also Fig. S1). The coverage is defined as the fraction of valid alignments.

### Evaluating Effect of Training Time and Size

We generated smaller datasets containing 50,000, 100,000, and 200,000 alignments. Next, we trained transformers on each of the datasets for 60 epochs with the original transformer training parameters. Of note, we started with random weights, and thus, no transfer learning affects these results. Next, we evaluated the performance of the transformers at the end of each epoch, with respect to the following metrics: (1) training loss, (2) validation loss, (3) fraction of invalid alignments, and (4) SC-error. The validation data contain 2,000 alignments (used to measure the validation loss), and the test data contain 1,000 alignments (used to measure the fraction of invalid alignments and SC-error). Fig. S8 shows the four metrics as function of the number of epochs. Note that the figures contain the four metrics together for comparing the correlation between the metrics. Each metric has a different range, and thus, there are multiple y-axes.

### Evaluating the Focus Learning

We generated a new dataset to measure the effect of the evolutionary parameters. We have used the same random topology and the same branch lengths as were used in PD14. Next, ten bins of each of the evolutionary parameters’ ranges were created: (1.0, 1.1), (1.1, 1.2) … (1.9, 2.0), and (0.000, 0.005), (0.005, 0.01) … (0.045, 0.05) for the A_I and A_D and R_I and R_D, respectively. To analyze the effect of A_I and A_D, for each of the A_I bins, all of the A_D were tested. In each case, 100 alignments were generated when the R_I and R_D values were sampled randomly from the range (0.00, 0.05), Fig. S9a (SPD11). The score of these alignments was averaged to create a total score for each bin. Of note, this resulted in 100 bins (ten different bins for A_I and ten different bins for A_D each with all of the others). Similarly, a dataset to analyze the effect of R_I and R_D was created (Fig. S9b). Next, we generated a dataset with high values of R_I and R_D. We specifically trained an “original” transformer on this dataset and compared its performance to the base transformer, i.e., the transformer that was not trained on the high R_I and R_D values.

Next, we generated three nucleotides’ datasets each one with a narrower range of indel parameters, i.e., A_I, A_D, R_D and R_I, branch lengths and root lengths (ND10, ND11, and ND12). We trained the transformers from the on the datasets starting with the dataset of the wider range (ND10). Then, the optimized transformers were used as the starting point for the next dataset (ND11). The optimized transformers from ND11 were the starting point of the next dataset (ND12). To evaluate the transformers performance, we used the test data of the narrower dataset (ND12).

### Generating Misspecification Dataset

To create misspecification datasets, we simulated alignments that have four different regions, each evolving under a different indel model, i.e., each part of the alignment has different indel parameters. The R_I and R_D are sampled from the ranges: [0.01, 0.02), [0.02, 0.03), [0.03, 0.04), [0.04, 0.05) for the first, second, third and fourth regions of the alignment, respectively. The A_I and A_D are sampled from the ranges: [1.01, 1.2), [1.2, 1.4), [1.4, 1.6), [1.6, 1.8) for the different regions, respectively. Those values were assumed both when simulating nucleotide and protein datasets. For protein sequences, each region was simulated assuming a different amino-acid replacement matrix: JTT (*43*), BLOSUM (*44*), mtART (*45*), and LG (*38*), for the first to the fourth domains, respectively. For nucleotide sequences, the GTR1, JC, GTR2, JC, were assumed for each region, respectively (*46*). The GTR1 and GTR2 frequencies are (0.37, 0.166, 0.307, 0.158) and (0.265, 0.182, 0.171, 0.382) for the frequency of T, C, A, and G, respectively. The assume entries in the rate matrix were: (0.444, 0.0843, 0.116, 0.107, 0.00027) and (1.387, 1.79, 0.431, 0.321, 4.947), for a, b, c, d, and e as stated in matrix Q (see Generating True MSAs Using SpartaABC section), for GTR1 and GTR2, respectively.

### Embedding of Multiple Sequence Alignments in High-Dimensional Space

The NLP approach presented here enables embedding an MSA in a high-dimensional space, i.e., it allows automatic features extraction, which could be utilized for downstream analysis. The high-dimensional vector is created within the translation process from a set of unaligned sequences. To obtain the embedded vector, the unaligned sequences were given as an input to the trained transformer. The vector is internally created by the encoder part of the transformer, and we have modified the code of the transformer to extract it (to reduce translation time, we skipped the decoder process).

The high-dimensional vector described above contains 1,024 times *n* entries, where *n* is the length of the input sentence, i.e., the representation of the unaligned sequences in one of the source languages (“concat” and “spaces” were used in this analysis for the source and target languages, respectively). Thus, this vector depends on the number of characters in the unaligned sequences. A representation of this vector, for three sequences of length up to five characters is given in Fig. S4.

For various downstream tasks, it is often desirable to compress this vector to a fixed size, i.e., a size which does not depend the sequence length (the compressed vector size does depend on the number of sequences). For the above example, the uncompressed vector is of size 1,024 x 15 and the size of the compressed vector is 1,024 x 5. Our compression process is exemplified on the above example for the three sequences of length up to five characters (Fig. S4). The first four entries of the first row are averaged and are assigned as the first entry of the compressed vector. The first four entries of the second row are averaged and are assigned to the second entry, etc. Thus, the first four columns are in fact compressed row wise to a vector of size 1,024. Of note, this size of 1,024 does not depend on the length of the unaligned first sequence. Let c1, c2, and c3 denote the three compressed vectors. Each pipe sign, “|”, forms a column of 1,024 characters. Since we have three unaligned sequences, we have two columns of the pipe sign. Let p1 and p2 be the vectors (each of size 1,024) that correspond to the pipe signs. The compressed vector is the concatenation of all vectors, in this order: c1, p1, c2, p2, and c3. One should emphasize that the pre-compressed representation already integrates information from all sequences due to the transformer self-attention mechanism, and consequently, the compressed representation also integrates information from all sequences.

Fig. S4a illustrates the embedding extracted from the last layer of the encoder part of the transformer. For each character in the unaligned sequences, i.e., the source sentence, there is a corresponding column of 1,024 entries. To create the compressed vector (Fig. S4b), the columns referring to characters of the first sequences are averaged row wise to create 1,024 entries, i.e., c1 in the main text. The fifth column, i.e., the column of the pipe sign, is column p1.

### Comparing Different Encoding and Transformers Architectures

We examined several source and target languages for the translation task. In addition to “concat” (see Main Text) we consider another possibility for encoding the input sequences, and we call the corresponding language “crisscross”. In this language, the first word is the first character of the first sequence and the second word is the first character of the second sequence. From the perspective of a transformer, it helps because it needs to learn closer rather than longer dependencies, compared to “concat”. As the input sequences are not necessarily of the same length, a gap character (“–”) is placed if no more characters are available for one of the sequences. For example, for the example input sequences in Fig. 1 (“AAG” and “ACGG”), the input sentence would be “A A A C G G - G”. Thus, the number of words is always twice the length of the longest sequence.

In addition to “spaces”, we additionally consider an alternative decoding for the output alignments, with the corresponding language called “pairs”. We note that each column of the output alignment can be one of the following: “AA”, “AC”, …, “A-”, “CA”, …, “T-”, “-A”, …, “-T”. In “pairs”, each such possibility is a word. Thus, for nucleotide sequences, the dictionary size is 24. For protein sequences, the dictionary size is 440. Note that “--” is an illegal word.

We encoded the same datasets with different representations: (1) “concat” and “pairs”; (2) “concat” and “spaces” (see Main Text); (3) “crisscross” and “spaces”. Then, we trained two transformers on each of the datasets, e.g., two transformers on the “concat” and “pairs”, and two different transformers on “concat” and “spaces”. The transformers were trained with the same tunable parameters. We compared the transformers’ performance on the same test data. This comparison was done on datasets ND3, PD2, and PD3 (see Table S3 in the Supplementary Text section). Note, that we did not use the “pairs” representations when shifting from pairwise to multiply alignment because the dictionary of pairwise and multiple alignments are different, which disenables transfer learning.

We also considered different architectures for the transformers, “BART” and “vaswani_wmt_en_de_big” (*14, 47*). Both types of transformers were trained with applying the “concat” with the “spaces” representations. We tested the results on proteins datasets: PD1, PD2, PD3, and PD4 (see Table S3). Of note, the transformers mentioned above are not pre-trained models and thus we start from random weights, and our own dictionaries.

### Evaluating the Transfer Learning

In transfer learning, the starting point of a transformer is another transformer that was already optimized on a specific dataset. We aimed to evaluate the contribution of transfer learning to the accuracy. To this end, we compared three different scenarios (illustrated in Fig. S5). In scenario 1, we evaluate a transformer that first encounters protein data PD5 (three protein sequences). This transformer was trained before on simpler datasets. In scenario 2, the trained transformer from scenario 1 was trained on additional more complex datasets (PD6, PD7, PD8, PD9, PD10, PD11, PD12, PD13, PD14, PD15) and was then re-trained on PD5. In scenario 3, the trained transformer from scenario 1 was retrained on PD5, without experiencing more complex datasets.

A similar evaluation was done on nucleotide transformers. Here instead of PD5, the base-dataset was ND4, comprised of alignments of three sequences. In scenario 2, the additional more complex datasets are: ND5, ND6, ND7, ND8, ND9, ND10, ND11, ND12, ND13, ND14. Figure S5 illustrates the three-stage training process of the transformers. For illustration purposes, the figure contains only five datasets. Consider “D3” to be the target dataset. In scenario 1 the transformer was trained on datasets “D1”, “D2” and “D3”. In scenario 2, the transformer from scenario 1, was then trained on more complex datasets: “D4” and “D5” and then retrained on “D3”. In scenario 3, the transformer from scenario 1, was retrained on “D3”, in order to reduce the difference to scenario 2.

### Extracting the Attention Maps

In the Fairseq library, which we used for training the transformers, there is an option to extract the attention map. Since using this option resulted in a bug, we fixed it by changing the source code of the library. Once the issue was fixed, we could save the attention maps.

### Comparing Against Other Alignment Programs

We compared BetaAlign against the following programs, used with default parameters: MUSCLE v3.8.1551 (*18*), MAFFT v7.475 (*19*), PRANK v.150803 (*20*), T-Coffee Version_12.00.7fb08c2 (*15*), ClustalW 2.1 (*16*), and DIALIGN dialign2-2 (*17*).

### Supplementary Text

#### Generalizing to Large MSAs

Aligning long sequences with transformers introduces new challenges. The attention map calculated inside the transformer is *O(n^2^m^2^)*, both in term of memory and running time, where n and m are the number of sequences and the sequences’ length, respectively. Note, the mn reflects the total words that need to be translated. Thus, increasing the alignment size causes memory and time problems. This is a known problem in the NLP domain, and so different efficient transformers were created in order to tackle this issue (*48*). Those architectures cannot be directly applied to the alignment problem (see the section on attention maps below). Hence, we developed a new way to align longer sequences by splitting the input sequences into short segments and training the transformers to align each segment (the size of the segments varies along the run, but is approximately 100 alignment columns, see Methods for details). We compared the performance of all aligners on MSAs of five sequences, with true alignment lengths up to 2,500 bases and 2,508 amino acids, for nucleotides and protein sequences, respectively. Clearly, the performance of BetaAlign is affected by the division of sequences into short segments, both when aligning nucleotide (Fig. S6a) and protein (Fig. S6b) sequences. Notably, even with this approximation, BetaAlign is more accurate than the widely used aligners MAFFT and ClustalW for nucleotide sequences and more than MAFFT, ClustalW, and MUSCLE for protein sequences.

#### Coverage Results

When generating the results of Fig. 2 in the main text, a small fraction of the resulting alignment was invalid, e.g., because of mutations introduced by the transformer to the unaligned sequences. As explained in the New Approach section, changing the tunable parameters of the transformers leads to different deep-learning models. We harnessed this feature of the neural network models and generated an additional transformer for aligning nucleotide sequences as well as an additional transformer for aligning protein sequences (we call each of these transformers “alternative”). When we aligned the invalid cases with the alternative transformers, a large fraction yielded valid alignments. Thus, every time the first “original” transformer fails, the results of the alternative transformer were considered. This methodology substantially increased the coverage, i.e., the percentage of MSAs that were successfully aligned. Specifically, the fraction of datasets that could not be aligned was less than 2% both for nucleotide and protein sequences (Fig. S7).

### The Effect of Training Time and Size

We next tested how the number of epochs and training size affect the accuracy and coverage of BetaAlign (in each epoch, the transformer processes the entire training data once, aiming to reduce the loss). We compared transformer performance trained on three training data sizes: 50,000, 100,000 and 200,000 protein alignments. Our results clearly indicate that for all datasets, the training loss decreases as the number of epochs increases, reaching almost a plateau when the data size is 200,000 alignments (Fig. S8). For each training data size, the validation loss follows the decrease in the training loss, suggesting that there is no over-fitting for the transformer. The coverage also continuously increases. For example, after 20 epochs the coverage was ∼ 40%, i.e., ∼ 60% of the resulting alignments were invalid. However, after 60 epochs, the coverage was already ∼80%. Of note, when studying the effect of training size, we started with naïve transformers, i.e., transformers that start with random weights. Hence, the coverage and accuracy are substantially lower than those presented in Fig. 2 (main text), and Fig. S2, in which we used transformers that were continuously improved using transfer learning.

The SC-error seems to substantially fluctuate even after 30 epochs (we note that the SC-error quantifies the error on valid alignments only, while the loss function quantifies the error on all alignments). The results further suggest that the loss function is correlated to the SC-error, but the correlation is mediocre at best. The correlation between the loss on the validation data and the SC-error on the dataset of 100,000 alignments, between epochs 20 and 60 was R^2^ = 0.467 (*P* = 0.0023). We speculate that this low correlation reflects the fact that the loss function is different from the SC-score (see “Additional Comments”).

Comparing the training and validation loss between the different training size datasets indicates that increasing the training size decreases the loss (training loss at epoch 60: 0.989, 0.985, 0.977, for datasets of 50,000, 100,000, 200,000, respectively). This gain in accuracy as reflected in the loss function was not evident when the performance is measured by the SC-error, probably reflecting the mediocre correlation between the two scores discussed above.

### The Effect of Indel Model Parameters on BetaAlign Performance

We studied the effect of the different parameters of the assumed indel model that generated the simulated data on the performance. To this end, we divided the alignments into bins by their evolutionary parameters: the insertion and deletion rate parameters (R_I and R_D, respectively) and the parameters that determine the distribution of indel lengths (A_I and A_D for the insertion and deletion distributions, respectively). As expected, increasing the indel rate parameters R_I and R_D substantially decreases accuracy (Fig. S9). The size of the indels has little effect on accuracy.

### Focus Learning

As stated in the New Approach section, we can train a transformer on a set of MSAs that share specific features, e.g., training them on MSAs with a high deletion rate and a low insertion rate. To determine if such a focus learning approach increases accuracy, we simulated three datasets. The first dataset (ND10) was simulated with a wide range of indel model parameters. The second dataset (ND11) was simulated on a sub-space of the indel model parameter space, i.e., the generated MSAs resemble each other in terms of indel dynamics. Finally, the third dataset (ND12) is even more restrictive in terms of the allowed indel dynamics (see Methods for details). Accuracy and coverage were measured on nucleotide datasets of five sequences with two different transformers (original and alternative). Our results clearly show that focus learning can improve both coverage and accuracy (Fig. S10), with the effect more substantial on coverage.

To further demonstrate the benefit of focus learning, we generated a dataset with high R_I and R_D (PD18) and trained the transformer. By using focused learning, we were able to decrease the SC-error from 0.029 to 0.025 and increase the coverage from 0.43 to 0.92. These results clearly show that focus learning has the potential to improve the accuracy of BetaAlign, and it is possible to generate different transformers for different regions of the space, which would ideally lead to a set of algorithms that outperform all other aligners across the entire space.

### Model Misspecification

In all the above analyses all positions within a specific MSA were generated by a model controlled by a specific set of parameters, in other words, there was no spatial model variation. We next evaluated how sensitive is BetaAlign to model misspecification. Specifically, we trained BetaAlign on alignments without spatial variations, but tested its performance on alignments generated with spatial variations. Test datasets were created by generating four “domains”, each with a different indel dynamics (see Methods). BetaAlign was substantially more accurate on the test protein dataset compared to other aligners (Fig. S11a). On the nucleotide dataset, BetaAlign SC-error was lower than two aligners but higher than the other two (Fig. S11b). We note that without model misspecification, BetaAlign had the highest accuracy compared to all other aligners on nucleotide datasets and was not the best for protein datasets. Model misspecification changed this order, as BetaAlign is now the most accurate aligner for protein dataset and had intermediate accuracy on nucleotide dataset. In general, machine-learning models are sensitive to deviations between the training and testing distributions. We found it surprising as in this case, the classic aligners on the protein dataset were more sensitive than the BetaAlign.

### Embedding Extraction for Downstream Tasks

The transformer is composed of two parts, the encoder and the decoder (see section New Approaches). After encoding the input sequences to a sentence, the encoder creates high dimensional vector representations of the source sentence, which are passed to the decoder to create the translated sentence (in our case, the aligned sequences). These vectors embed the information in the unaligned sequences, i.e., they are a numeric representation of the information stored in the set of unaligned sequences. We compressed these vectors to a vector of fixed size. In the case of five sequences, the number of features is 9,216 (see Methods).

To exemplify the usefulness of such a representation, we used the resulting features for a different machine-learning task, which is to estimate the length of the sequences of the root of the tree that generated the resulting sequences. To this end, we trained a linear regression model on these features, based on a training set of 90,000 nucleotide MSAs, each with five sequences (ND10). The accuracy of the linear-regression model using these features was evaluated on test data comprising 10,000 MSAs. Fig. S12 demonstrates the correlation between the true and inferred root lengths. A very high *R*^2^ of 0.91 with a *MSE* of 2.003 was obtained, suggesting that our approach can be used to compactly code sequences, as a preliminary step for downstream machine-learning tasks.

### Testing Alternative Encoding Schemes

Table S1 displays the performance of transformers trained with various encodings for pairwise protein alignment (as explained in New Approaches). All six variants had similar accuracy, as measured by the SC-score. The “pairs” target language resulted in the highest coverage. For the task of multiple sequence alignments, we decided not to work with the generalization of the “pairs” target language because in this case the size of the dictionary increases exponentially with the number of input sequences (if we denote n as the number of input sequences, the output protein dictionary size is 22^n^ − 1). Furthermore, because the dictionary size depends on *n*, the “pairs” target language hinders using transfer learning. Note that the “spaces” and “crisscross” dictionaries do not change when increasing the number of input sequences. As the performance using the “concat” source language was slightly better than that of the “crisscross” source language, the former was applied for all analyses.

### Architecture Comparisons

We compared the results of two transformers architectures: “BART” and “vaswani_wmt_en_de_big”. While both “BART” and “vaswani_wmt_en_de_big” architectures contain 16 attention heads, with an embedding size of 1,024, they differ in the details of their network design, including a different number of layers: 6 and 12 for “vaswani_wmt_en_de_big” and “BART”, respectively (*14, 47*). The performance of these architectures, tested on four different datasets of amino-acid sequences, and with several different sets of internal parameters (max tokens and learning rate), clearly shows that both the coverage and the SC-score are higher for the “vaswani_wmt_en_de_big” architecture (Table S2). We thus selected this architecture for all further analyses.

### Transfer Learning

Our approach heavily depends on transfer learning. We use it both to increase the complexity of the datasets and to focus transformers into a subregion of the evolutionary parameters range. Except for the first datasets (where the weights are randomly initialized), all the transformers are based on the previous dataset. The transformers of the nucleotide datasets have a different path of training from the transformers of the amino-acid datasets. In addition, the original transformer is based on the previous original while the alternative transformer is based on the previous alternative transformer, etc.

To measure the effectiveness of the transfer learning feature, we reran datasets and measured the SC-score. This was tested on two datasets of protein (PD5), and nucleotides (ND4) with lower SC-score compared to different aligners. As those datasets were at the start of the transfer path, the base transformers were trained only on millions of true alignments. We tested three alternative scenarios (detailed in the Supplementary Text). Briefly, the transformer in scenario 1 (transformer 1) is trained once on the target dataset. Transformer 2 (scenario 2) starts from the end point of transformer 1. It was then trained on various other datasets, and then re-trained on the same target data (either PD5 or ND4). Transformer 3 also starts from the end point of transformer 1, but is only retrained on the same target data without experiencing more complex datasets in its path. Transformer 2 outperformed transformer 3, which outperformed transformer 1, clearly demonstrating the benefit of transfer learning (Fig. S13).

One could also compare the performance demonstrated at Fig. S8 to the performance obtained in Fig. 2. In Fig. S8, the transformer was trained on a random starting point resulting in a fraction of invalid alignment of 0.21 and an SC-error of 0.03. The transformers of Fig. 2 were trained on optimized starting point resulting in a fraction of invalid alignment of 0.006 and an SC-error of 0.0245. These results demonstrate that the transfer learning boosts the performance of BetaAlign. As seen from Fig. S8, additional improvement is not expected when increasing the training time.

### Attention Map Comparison

BetaAlign harnesses NLP-based translation methods to analyze biological data. However, it is clear that biological data are substantially different from natural languages data, e.g., they have a much smaller dictionary size (six for nucleotides using the “spaces” representation, compared to an order of a hundred thousand words in vocabularies of natural languages). Attention maps provide a gateway for understanding the internal properties of various transformers (explained more below). Put simply, attention maps show on which part of the sequences the transformer focuses while generating the next word. Fig. S14a displays the averaged attention map gathered from the attention layers of the decoder (in this case, when processing an amino-acid MSA of seven sequences, from PD14). Fig. S14b shows an attention map from an NLP English to French translation task (*49, 50*).

Note that each attention map corresponds to translation of a single sentence. In the case of Fig. S14b, the translation is of the sentence “The agreement of the European Economic Area was signed in August 1992.”. The sentence translated in Fig. S14a is a concatenation of seven unaligned sequences of length ∼ 40 nucleotides. Clearly, natural language sentences are much shorter, on average, than sentences translated by BetaAlign, and thus, the BetaAlign attention map is much larger (consisting of hundreds of words) than the natural language one (consisting of dozens of words). Moreover, the attention in the natural languages is focused on one diagonal while the attention in BetaAlign is focused on several diagonals, the number of which equals the number of inputs sequences. The difference stems from the fact that in the BetaAlign translation task, generating the next word requires assigning more weight to information that is distributed among all sequences. Of note, this analysis was conducted on the “concat” and “spaces” for the source and target languages, respectively.

Fig. S14c is an enlargement of the attention map around the rightmost diagonal of Fig. S14a. The x-axis refers to the characters of the input sentence and the y-axis refers to the translated characters of the output sentence. As can be seen, the diagonal is actually a step of seven rows and one column. This is the logical explanation as each of the seven rows extracts information from this column, i.e., the characters of a column at the resulting MSA are all dependent on the same character in the unaligned sequences. The red arrows mark the corresponding output and input characters. As can be seen, the weight is higher in this area, i.e., the transformer learned the importance of the corresponding area.

Due to the dynamics of our attention maps (specifically, the focus split), we cannot easily use NLP architectures for longer sentences, which have dedicated attention maps (*48*). In our segmentation strategy, instead of calculating the entire attention map, we focus the attention on specific places of the diagonals. Future research will focus on improving attention calculations.

### Runtime

The training phase depends on the size of training data and the number of epochs, i.e., how many cycles of processing the entire training data were executed. The typical training time of an epoch on a cluster of eight GPUs is approximately 25 minutes for a dataset of 100,000 alignments comprised of three sequences with an ancestral root length with an average of ∼ 40 amino acids. The total number of epochs needed to reach convergence of the loss function depends on the starting point of the transformers used. Starting from random weights will take substantially more time than a transformer that was initiated by transfer learning.

Once the transformer is trained, aligning a set of up to ten sequences of similar lengths to the training data takes a few seconds on a GPU. However, we note that the time complexity of BetaAlign is quadratic for processing short sequences, but linear to the number of sequences when using “segmented transformers”, which should provide an advantage for BetaAlign in terms of running times when analyzing MSAs with a high number of long sequences.

### The Effect of Indel Model Parameters on Other Aligners

The following graphs display the effect of the indel model parameters on the different aligners (Fig. S15). Similarly, to the results observed in “The effect of Indel Model Parameters on Performance”, the R_I and R_D have substantially higher effect on the performance than the A_I and A_D. One can see that different aligners respond differently to the indel dynamics.

### Additional Comments

1. As each segment translated with the “segmented transformers” is translated independently, different alignments dynamics could be inferred based on this specific segment. This could have an impact on aligning large genomic sequences as different regions in the genome are corresponding to different dynamics as selection powers work differently across the genome.
2. In this research, we used classical NLP architectures, and loss functions. As we showed in the in Attention Maps Comparison section, biological data behave differently from conventional NLP translation tasks, which suggests that there is much room for improvement. Future research should aim to develop NLP tools that better capture the structure of biological macromolecules and their evolution. Specifically, we anticipate the future development of novel architectures of transformers as well as more relevant loss functions.
3. One of the main challenges of BetaAlign is generalizing the alignments for longer sequences, a well-known hurdle in NLP research. Here, we presented a way to divide the alignments into segments. Further work may explore alternative solutions to this challenge. Different architectures could increase the length of the sentences, and different representations could reduce the number of words in each sentence, e.g., by compressing both the input and output sentences.

### Additional Datasets

Table S3 contains the parameters used to generate the different datasets. (a) contains the information on the DNA datasets; (b) contains the information on the protein datasets; (c) contains information about other specific datasets. The random topologies were generated using the program ETE version 3.0. For each dataset, millions of alignments were generated and divided to training data, validation data, and testing data.

### Data for Specific Figures

The datasets for Fig. S6 are SND5 and SPD9 for the nucleotide and protein dataset, respectively. To create Fig. S7, we used the same dataset as Fig. 2 in the main text. To create Fig. S8, we generated smaller datasets containing 50,000, 100,000, 200,000 alignments sampled from the data created for Fig. 2 (main text). Of note, we have used the three input sequences of protein (SPD1). To create Fig. S9, we generated a specific dataset SPD11, with specific alignments for each one of the evolutionary bins (see “Evaluating the Focus Learning”). The transformers of Fig. S10 were trained on datasets ND10, ND11, and ND12 and were evaluated on the test data of ND12. In Fig. S11, the transformers that we used were trained on datasets SND1 and SPD1. The test datasets that include spatial variation are SND6 and SPD10 for nucleotide and protein datasets, respectively. For Fig. S12, we encoded 100,000 sentences representing the unaligned sequences and extracted the vector from the last layer of the encoder. To this end, we used a dataset composed of five nucleotide sequences, ND10. The datasets for Fig. S13 are PD5 and ND4 for the protein and nucleotides datasets, respectively. Shown in Fig. S14a and Fig. S14c are attention maps of different resolutions extracted from one of the protein alignments composed of seven sequences (dataset PD14).

---

**Fig. S1.**
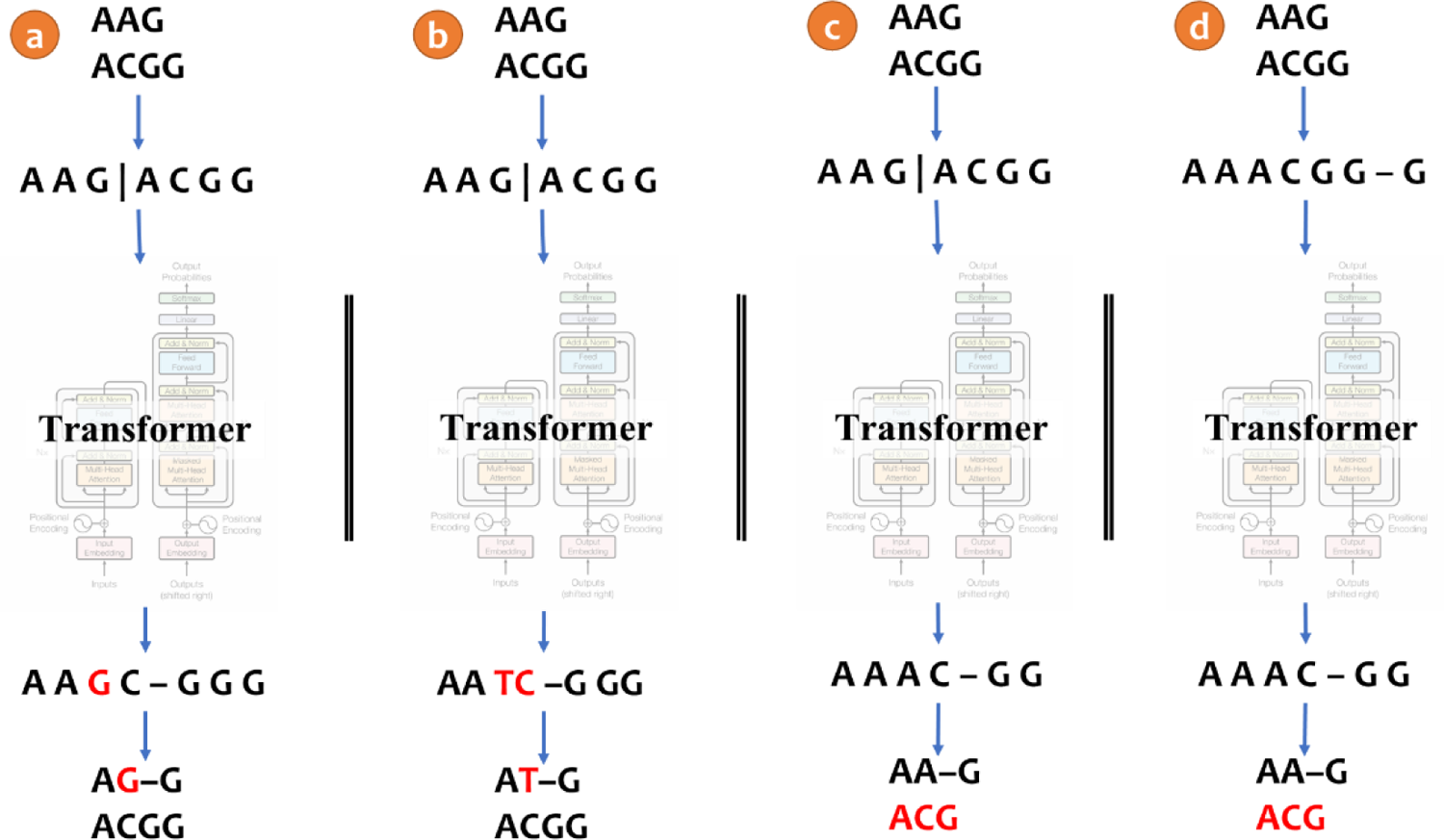
Invalid alignments. (a) Although the first input sequence is AAG, the first row in the resulting pairwise alignment is AG–G, which is invalid as the algorithm not only introduced gaps but also mutated a character. Here, sequences were encoded using the “concat” representation and decoded with “spaces”; (b) A similar problem may arise when decoding with “pairs”; (c) An example of an invalid pairwise alignment when applying the “concat” and “spaces” representations, stemming from an unequal length of rows in the resulting MSA. Note that this problem can never appear using “pairs”; (d) A similar problem may arise in the “crisscross” encoding. The transformer architecture illustration is adapted from (14).

**Fig. S2.**
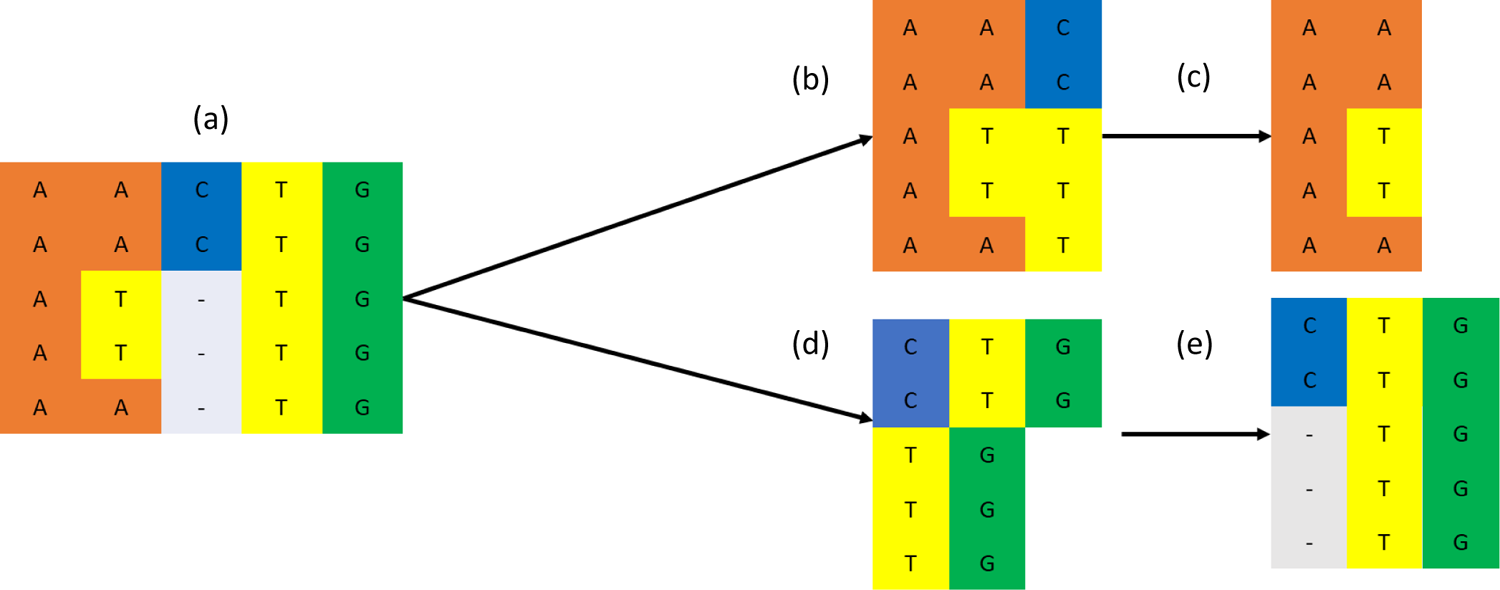
Creating training data for segmented transformers, explained using a segment size of three characters. (a) the true alignment; (b) The unaligned sequences of the first segment, i.e., the first x as described in the Methods section; (c) The segment of the true MSA that corresponds to the x presented in b, i.e., the first y. Note that the third column of the true MSA was not selected as it contains gaps, and the forth column was not selected, as it includes four characters of some of the original sequences (i.e., it is longer than the segment size of three); (d) The unaligned sequences of the second segment; (e) The corresponding segmented true alignment.

**Fig. S3.**
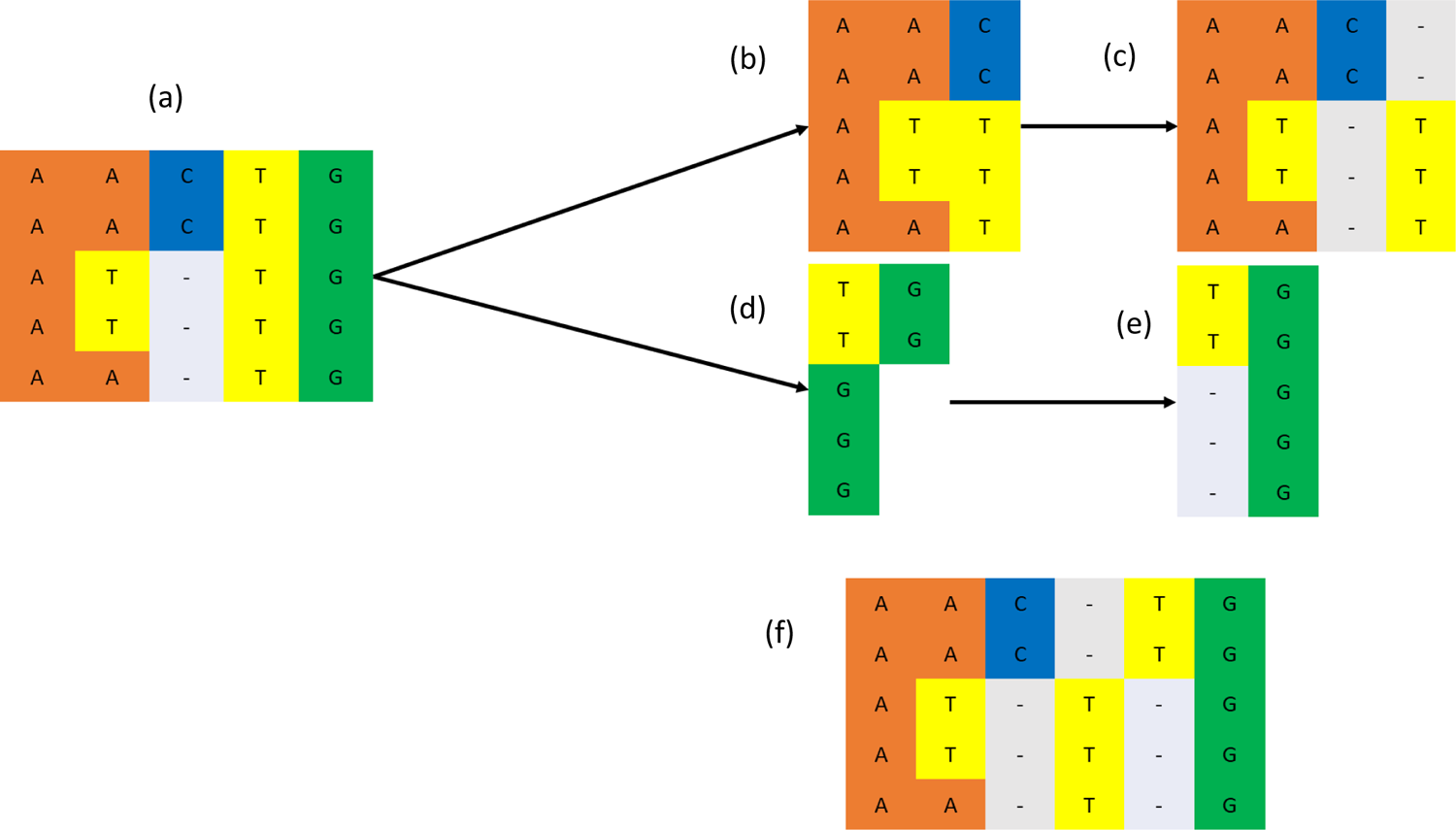
Importance of indel-free last column of segments. Consider (a) to be the true alignment, with a segment size of three characters. (b), (c) refers to the unaligned sequences and the aligned results, of the first segment respectively. Similarly, (d), (e) refer to the unaligned sequences and the aligned results of the second segment, respectively. (f) is the concatenation of the results. Note the difference between (a) and (f) as the third and fourth columns at (f) should be the fourth column in (a).

**Fig. S4.**
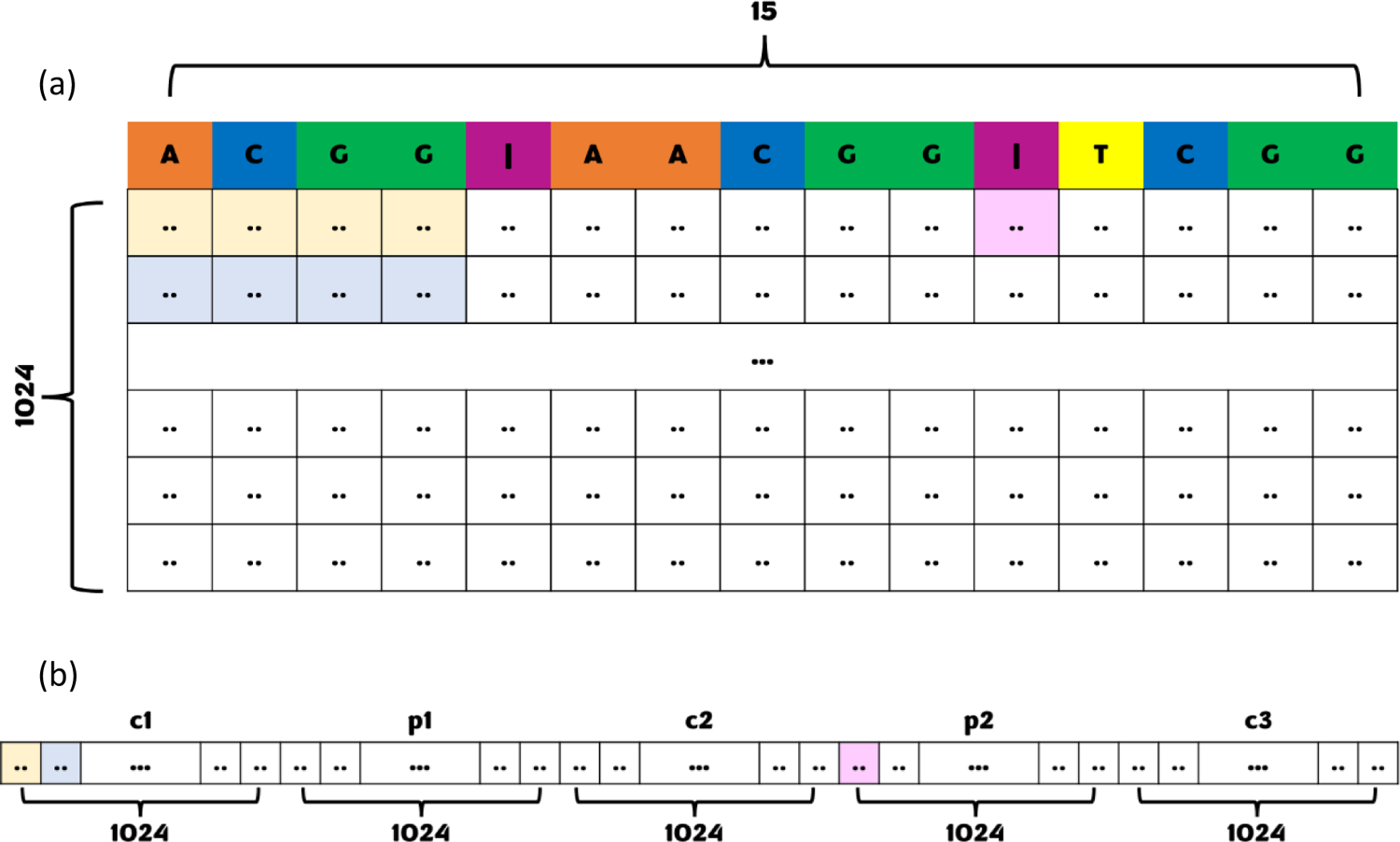
Example of compressing the embedding vector to a fixed size. Consider (a) to be the embedding of three sequences. The embedding dimension is of 1,024 x 15 as there are 15 characters is the source sentence. The compressed vector is (b) is of 1 x 5,120 as each one of the input sequences and the pipe sign correspond to 1,024 entries in the compressed vector. We have colored the first four entries of the first and second row, and the 11^th^ entry of the first row. The corresponding entries in the compressed vector are colored the same.

**Fig. S5.**
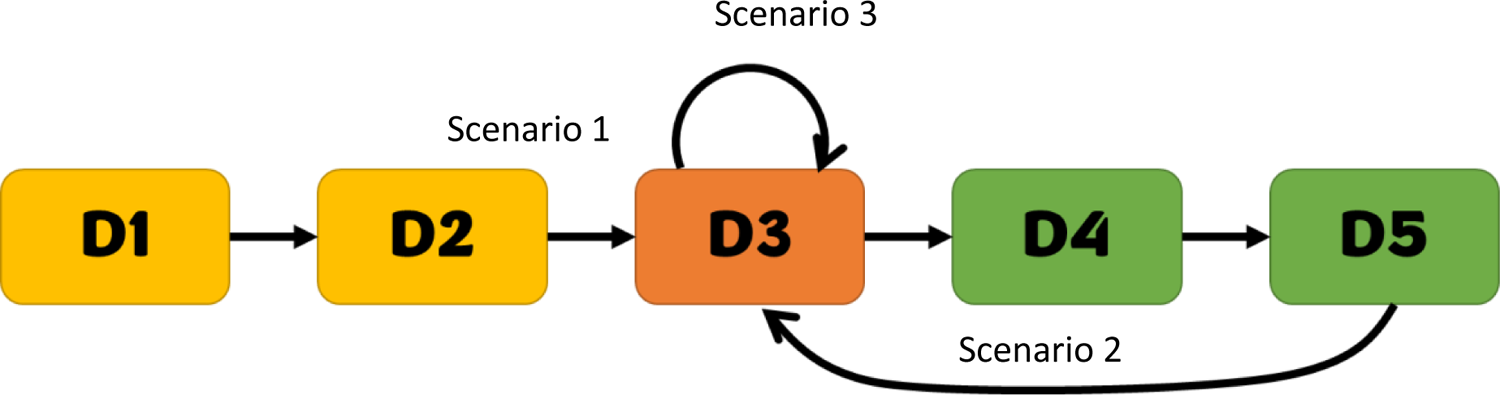
Illustration of the three scenarios to evaluate the transfer learning. Scenario 1 includes training on “D1”, “D2” and “D3”. Scenario 2 includes training on “D1”, “D2”, “D3”, “D4”, “D5” and “D3”. Scenario 3 includes training on “D1”, “D2”, “D3”, and “D3”. The datasets in yellow represent simpler datasets which were trained on before. The dataset in orange represents the target dataset, i.e., the dataset on which the performance was tested. The datasets in green represent more complex datasets that were trained on after the target dataset. Arrows between datasets represent the transfer learning, i.e., the transformer optimized on a dataset was used as a base transformer for the other dataset.

**Fig. S6.**
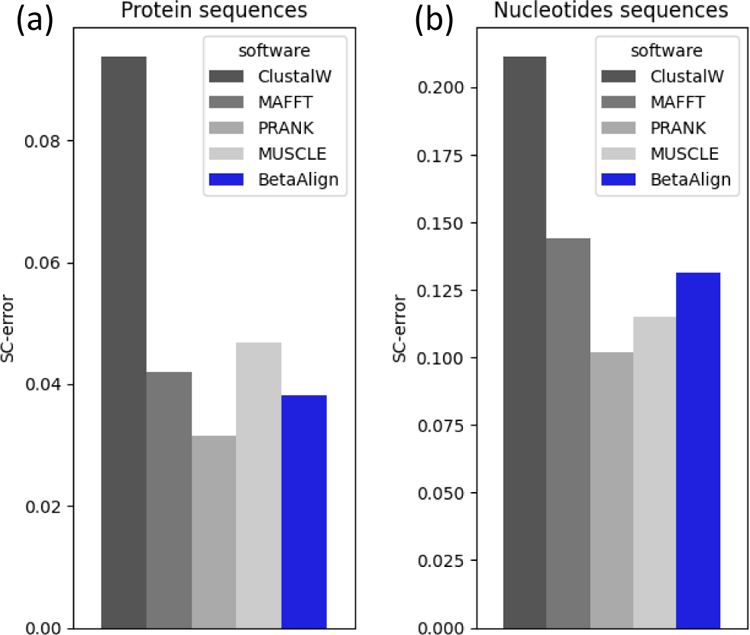
Running on large MSAs. Comparing the results of different aligners on longer sequences for five sequences for nucleotide (a) and protein (b) sequences. The average MSA length was 760 bases and 765 amino acids, for nucleotide and protein sequences, respectively. These datasets were aligned after dividing the input into short segments, thus overcoming computational memory and run time limitations introduced by large MSAs.

**Fig. S7.**
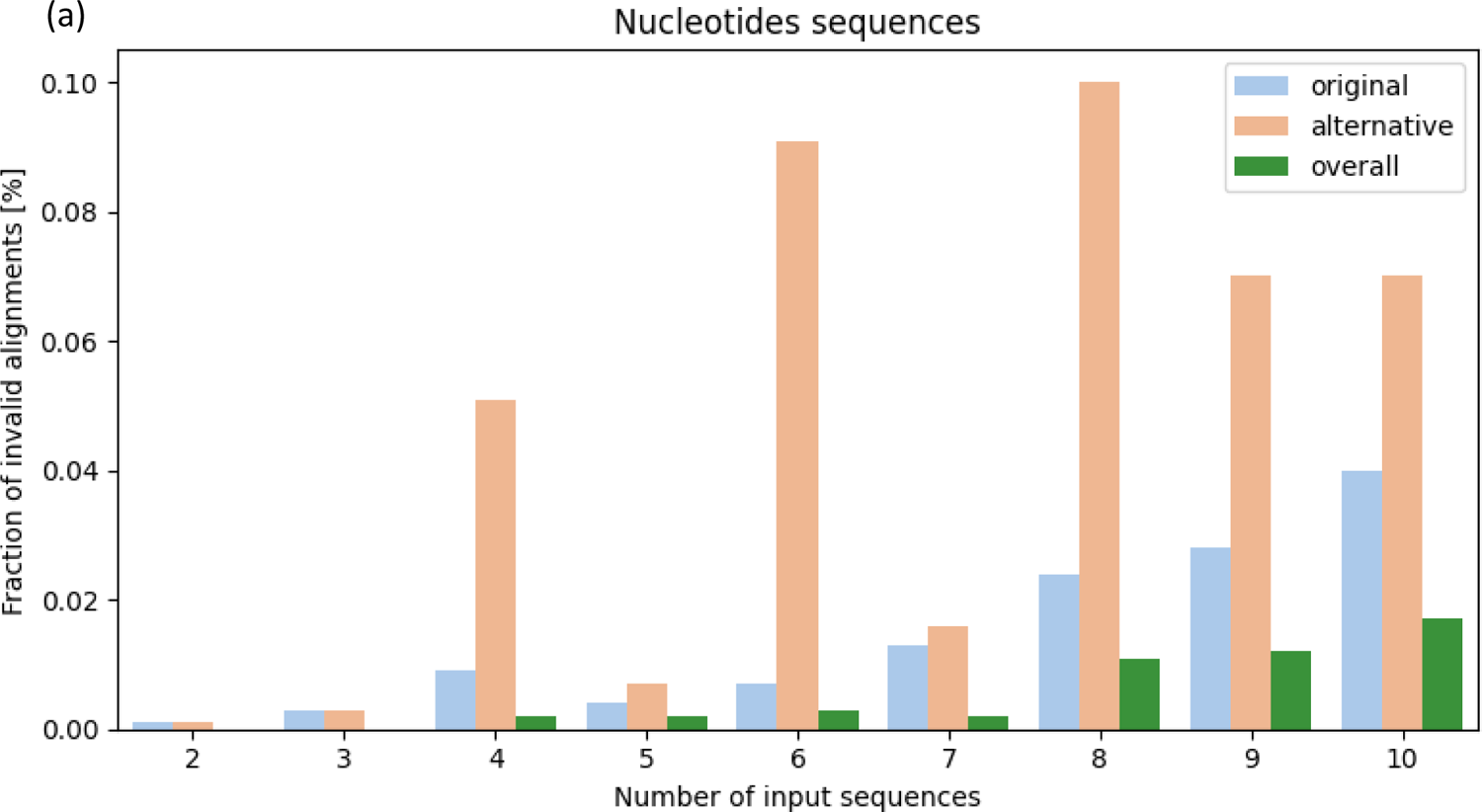
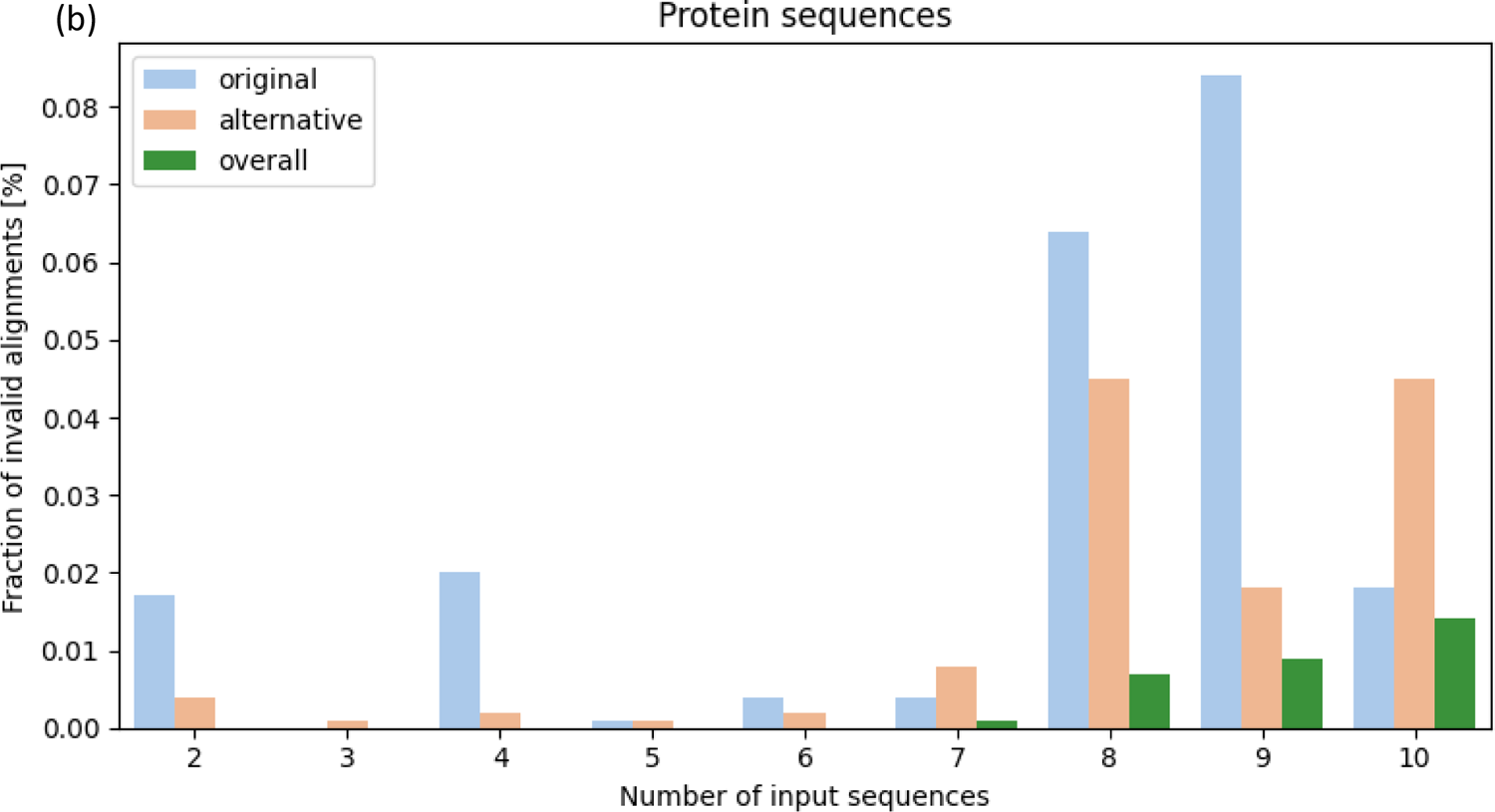
The fraction of invalid alignments for nucleotides (a) and protein (b) sequences as a function of the number of inputs sequences. For each dataset, two different transformers were trained. The fraction of invalid alignments of the “original” and “alternative” transformers are in blue and orange, respectively. In green is the fraction of invalid MSAs shared by both transformers.

**Fig. S8.**
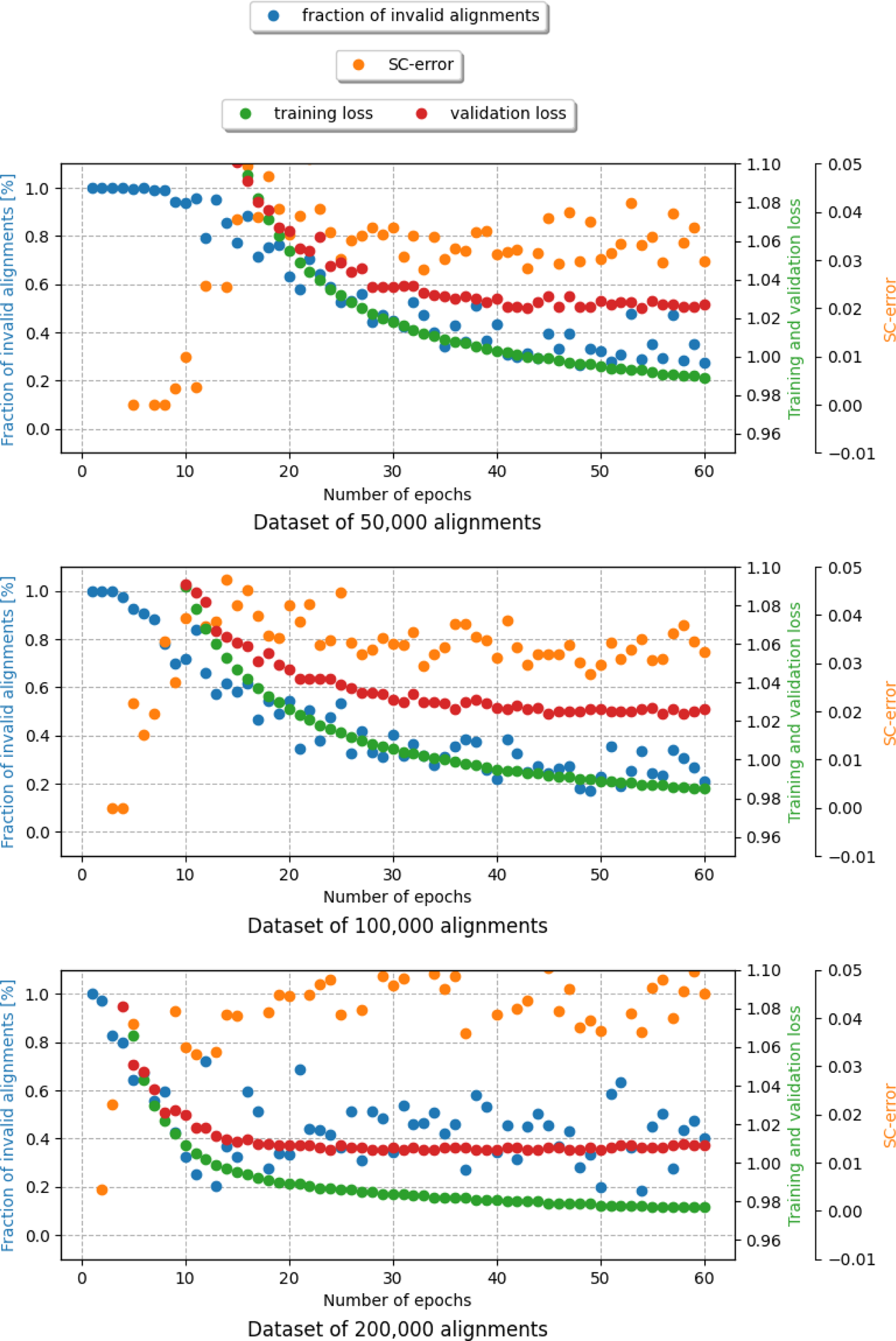
Effect of increasing the training time and size on the fraction of invalid alignments (blue dots), SC-error (orange dots), validation loss (red dots), and training loss (green dots). All alignments were of three sequences, dataset SPD1.

**Fig. S9.**
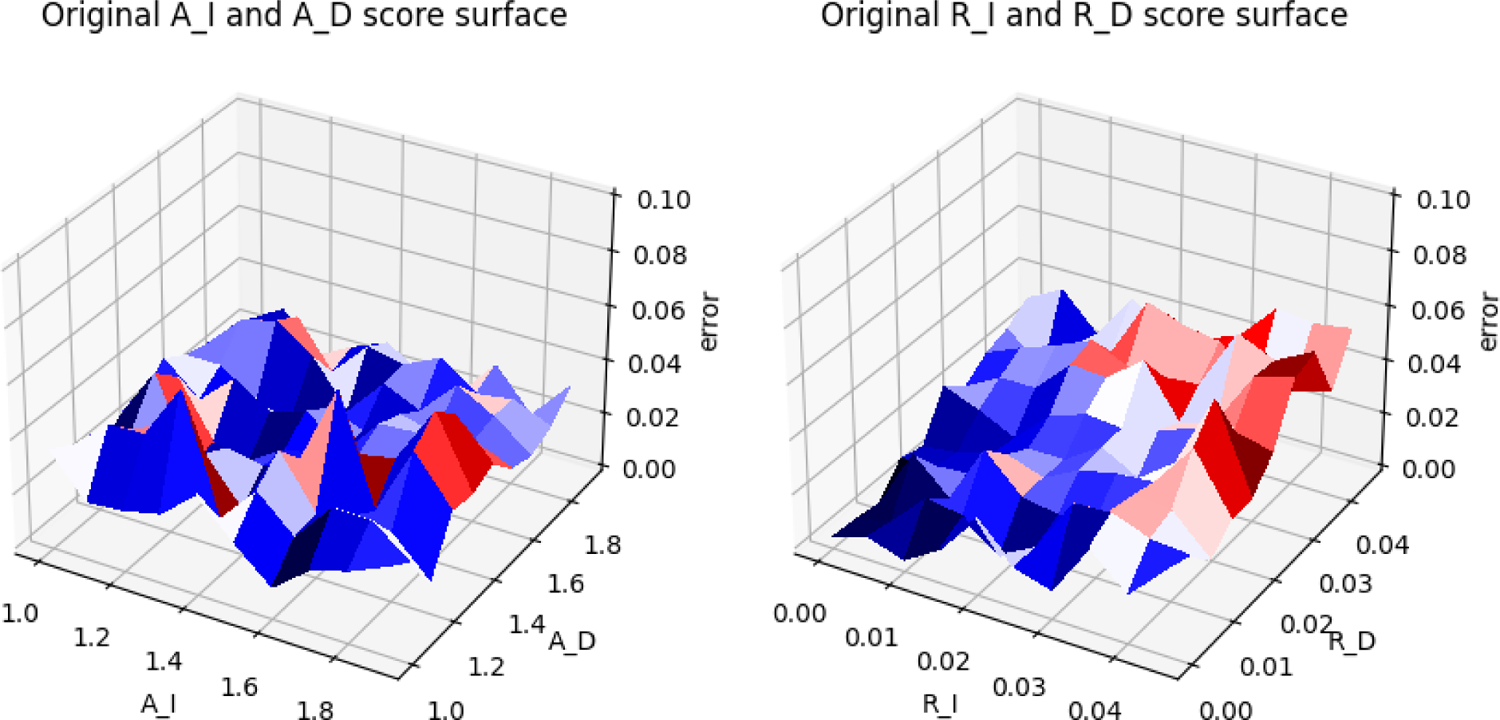
The effect of the indel model parameters on BetaAlign performance: (a) measuring the effect of A_I and A_D (in this case R_I and R_D were sampled from the entire range); (b) measuring the effect of R_I and R_D (in this case A_I and A_D were sampled from the entire range).

**Fig. S10.**
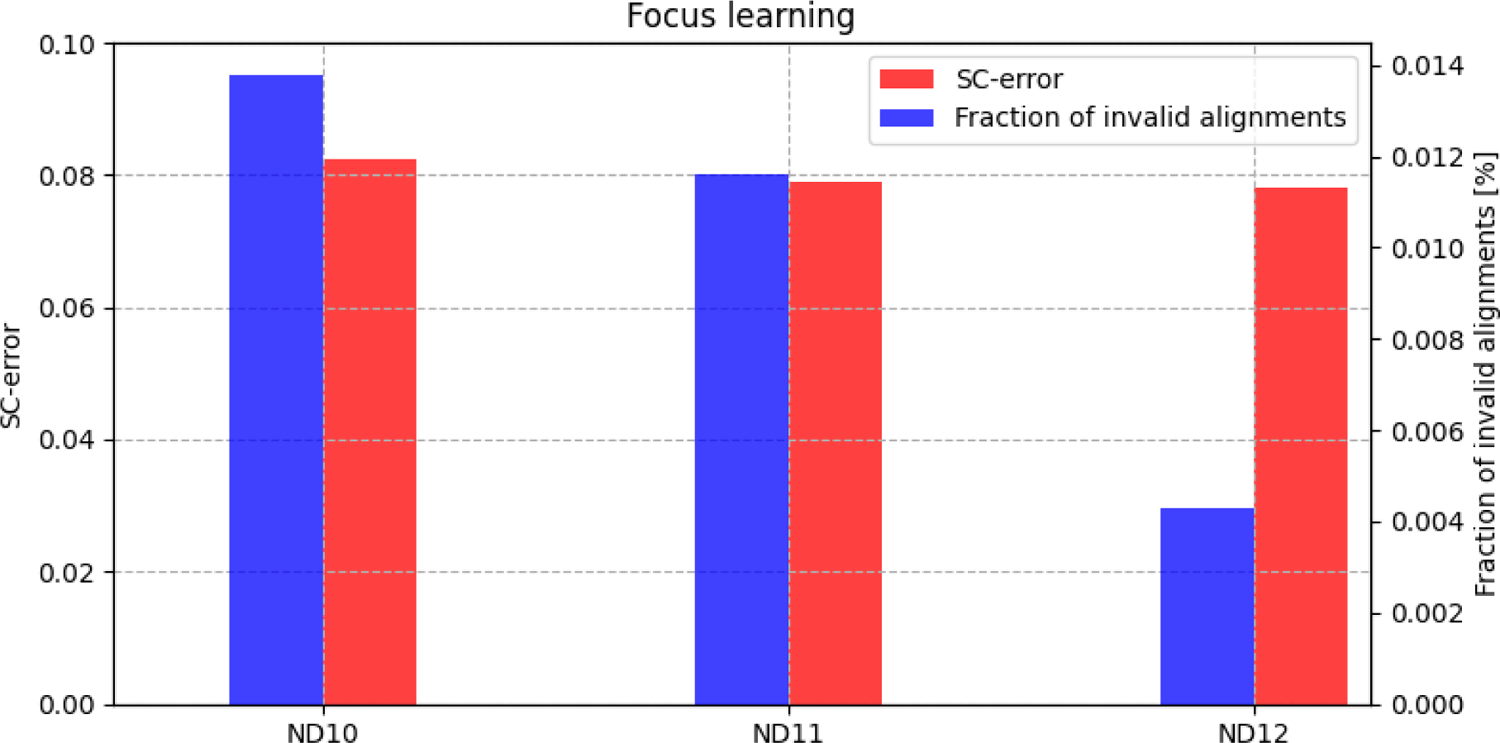
Effect of focus learning on the fraction of invalid alignments (a) and the SC-error (b). As the dataset number increases, the range of the indel parameters model decreases. Each dataset is a subset of the previous dataset, i.e., it was generated with a narrower range of indel model parameters.

**Fig. S11.**
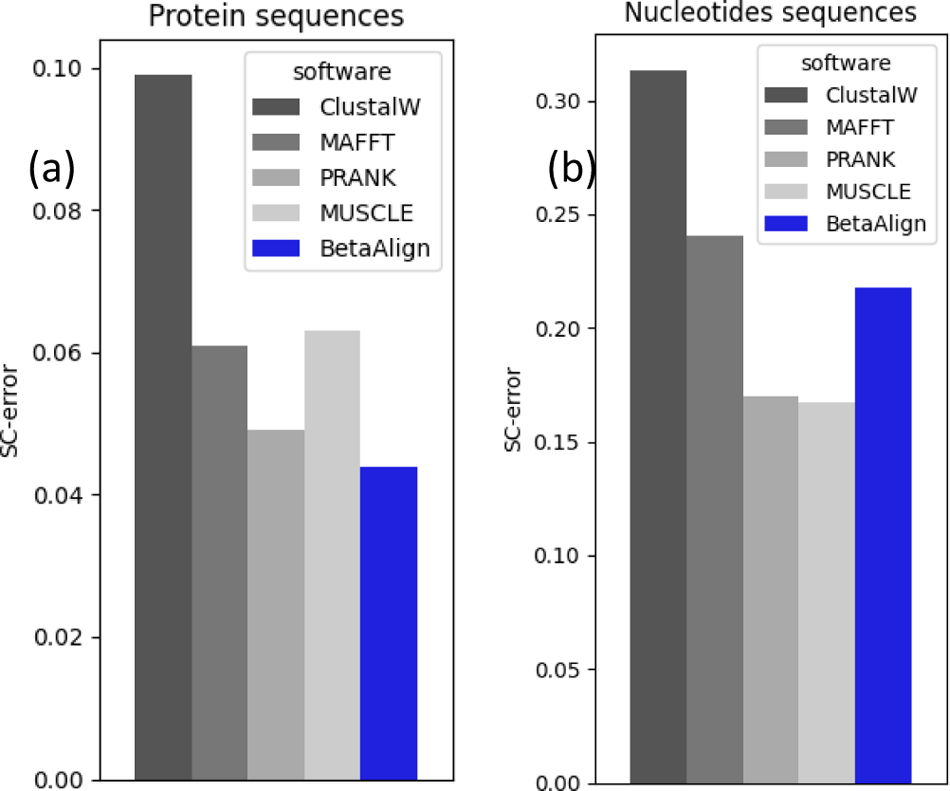
The effect of model misspecification on the accuracy of protein (a) and nucleotide (b) datasets. Train and test alignments had 10 sequences. Spatial variation was only introduced to the test data, i.e., different regions of the alignment were simulated with different sets of indel model parameters.

**Fig. S12.**
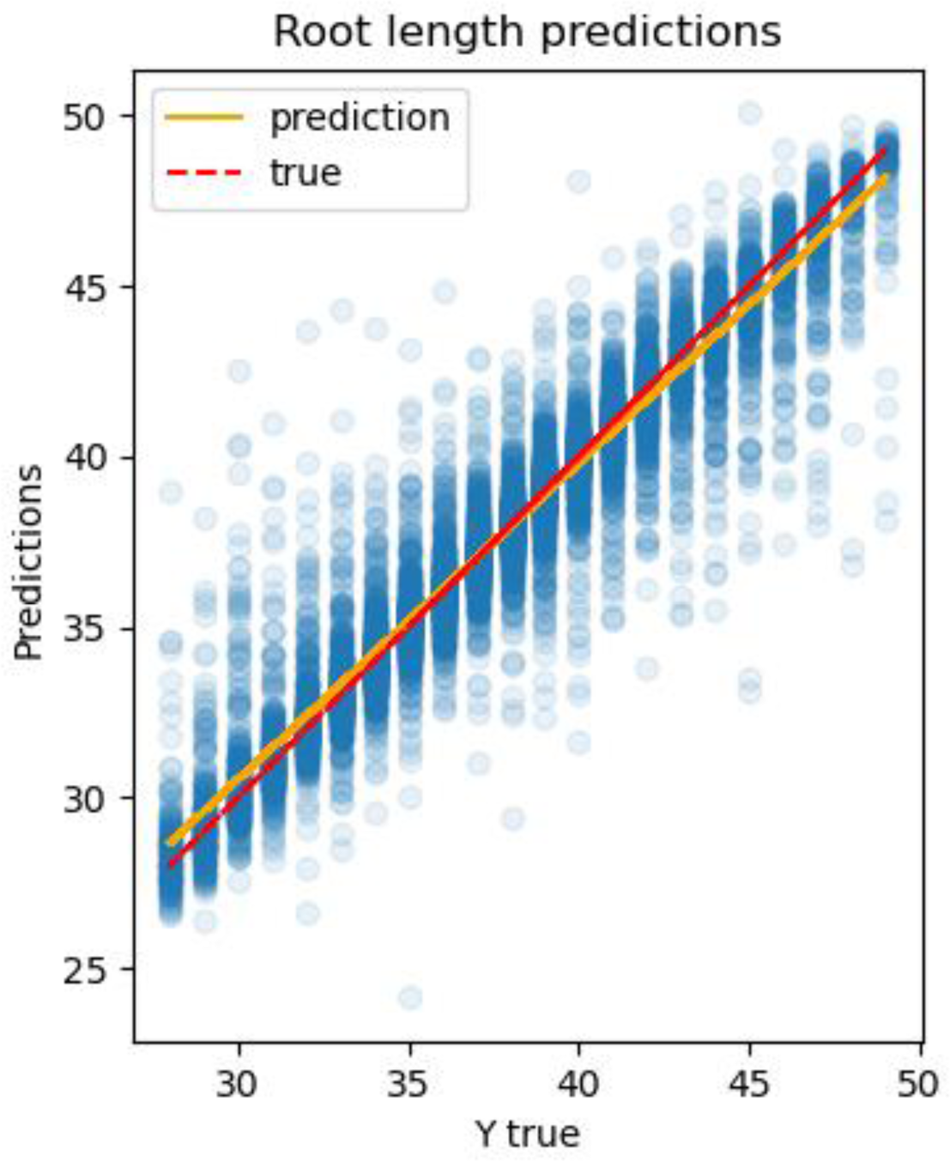
Results of the linear regressor we trained to predict the root length from the embedding of the unaligned sequences, with an R^2^ of 0.91 and *MSE* of 2.003. The orange line is the regression line, and the red line reflects the Y = x function.

**Fig. S13.**
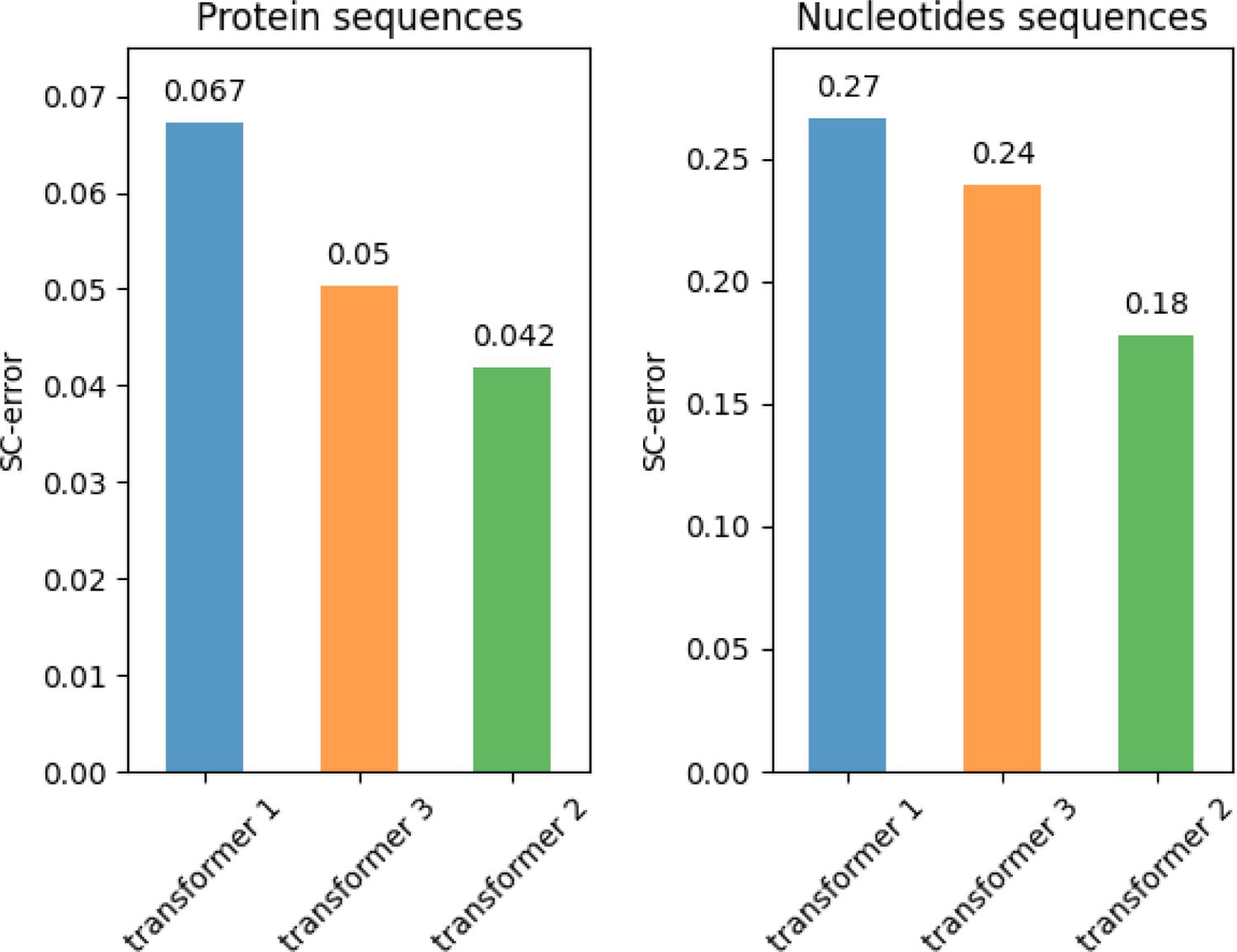
Effect of transfer learning on the SC-score reduced the error by 37.6% and 33.4%. We tested it on an amino acid dataset and nucleotides dataset, both containing three sequences.

**Fig. S14.**
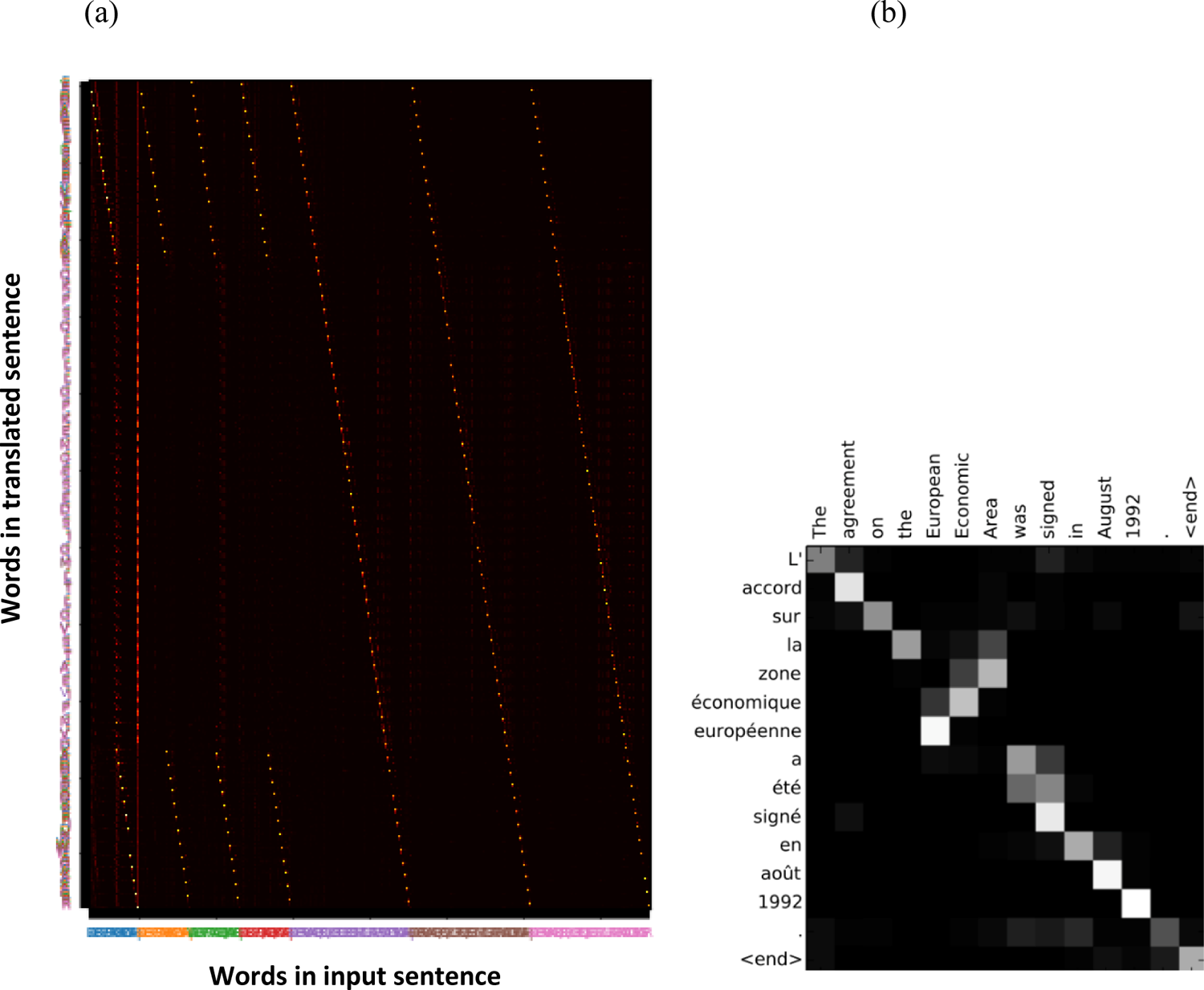
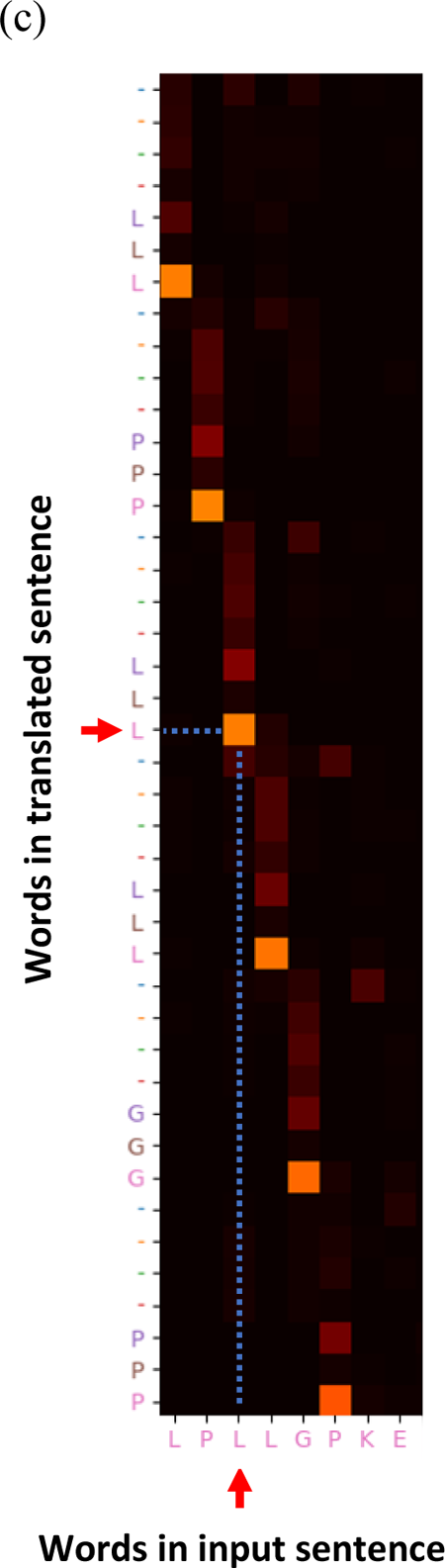
Comparing the attention maps of BetaAlign and other NLP tasks. The x-axis is the input words, and the y-axis is the output words. The attention splits between all the input words when generating the next word. (a) attention map extracted from the last layer of the BetaAlign’s decoder. One can see that the NLP task (b) refers to only one location of the input sentence while BetaAlign refers to multiply locations, one for each of the input sequences. To visualize this, we colored each of the unaligned sequences in a different color as well as the amino acid that refers to this unaligned sequence. When zooming into the map, there is a step of the attention of the four left diagonals, that we identified as an indel. In this indel, the first “|” have a meaningful part. (c) zooming in to one of the diagonals of (a). The red arrows represent the corresponding output and input characters “L”.

**Fig. S15.**
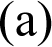
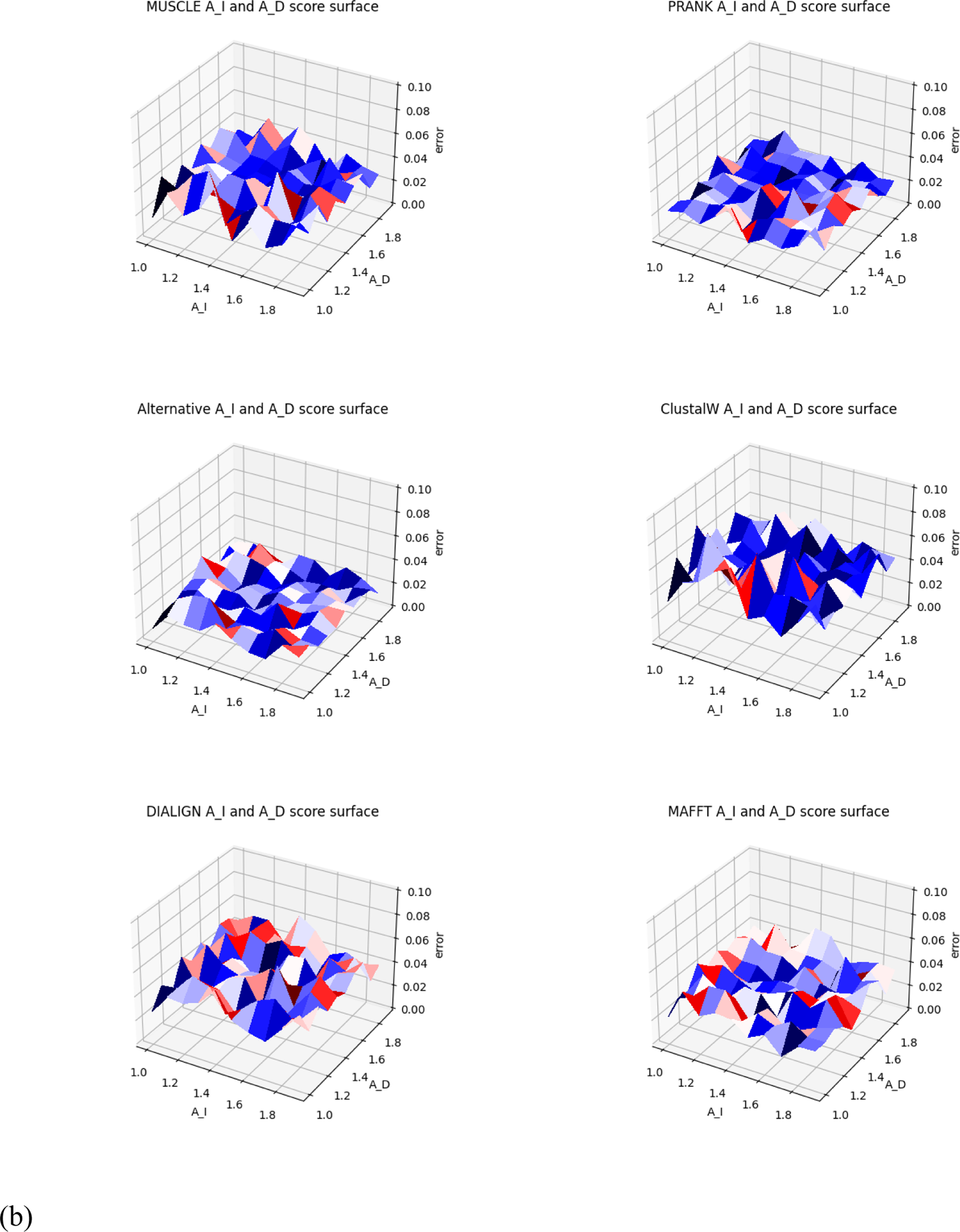
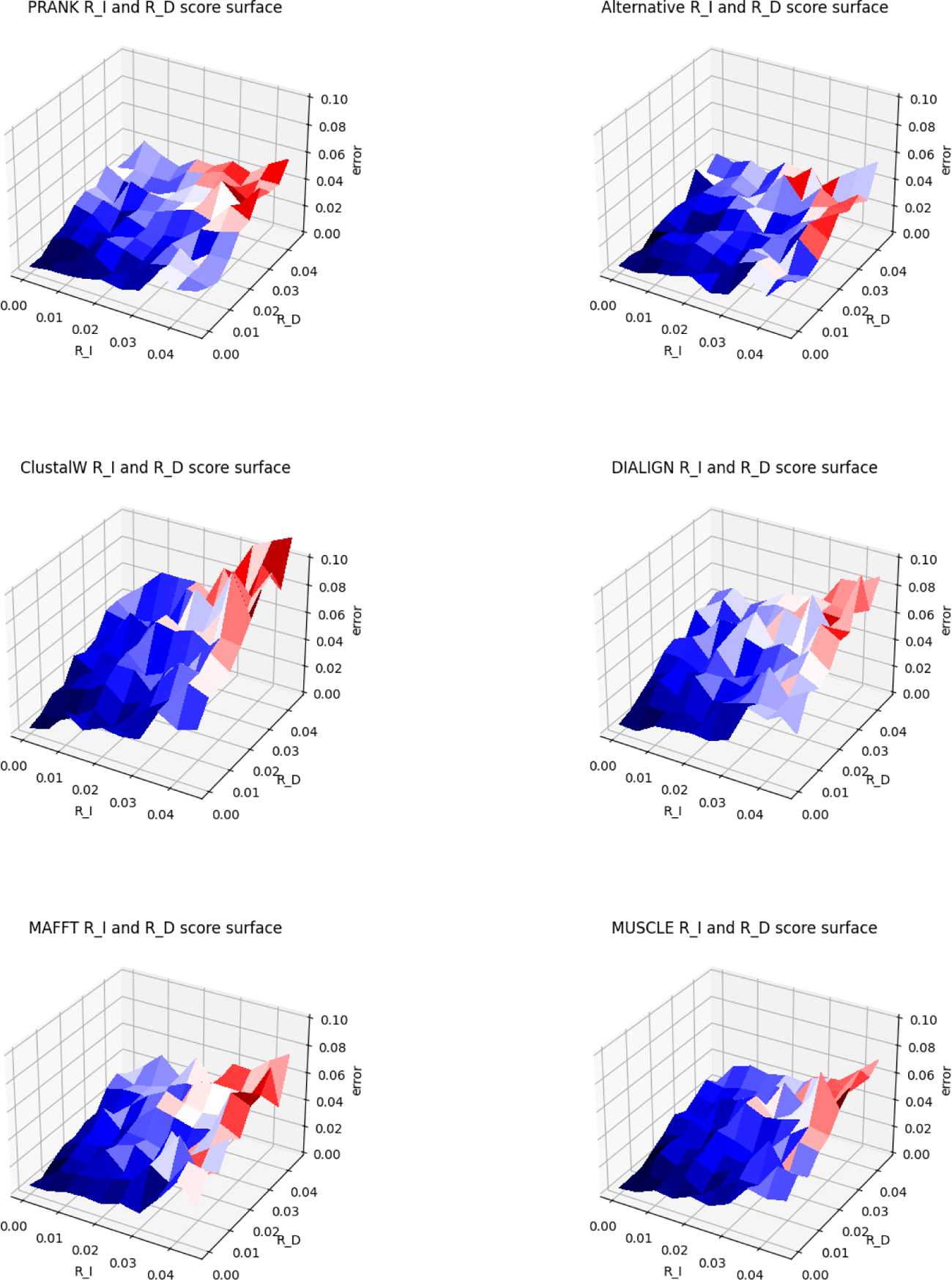
The effect of the indel model parameters on different aligners performance: (a) measuring the effect of A_I and A_D (in this case R_I and R_D were sampled from the entire range); (b) measuring the effect of R_I and R_D (in this case A_I and A_D were sampled from the entire range).

**Table S1.** Transformers performance trained on pairwise protein alignments with source and target languages (dataset PD2). Each combination of source and target language was tested with two values for the max-token parameter, resulting in two transformers: “original” and “alternative”.

| *transformer name* | *source language* | *target language* | *max tokens* | *SC-score* | *Coverage* |
| --- | --- | --- | --- | --- | --- |
| original | concat | *pairs* | 4096 | 0.997 | 0.939 |
| alternative | concat | *pairs* | 2048 | 0.997 | 0.952 |
| original | concat | *spaces* | 4096 | 0.997 | 0.835 |
| alternative | concat | *spaces* | 2048 | 0.998 | 0.635 |
| original | crisscross | *spaces* | 4096 | 0.991 | 0.503 |
| alternative | crisscross | *spaces* | 2048 | 0.995 | 0.636 |

**Table S2.** We trained both transformer architectures on the same amino-acid datasets and measured the accuracy and coverage. Those datasets were encoded with the “concat” as a source language and “spaces” as the target language, datasets PD1, PD2, PD3 are of pairwise alignments and dataset PD4 is of 3-MSA alignments.

| *transformer name* | *architecture* | *max tokens* | *learning rate* | *SC-score* | *Coverage* |
| --- | --- | --- | --- | --- | --- |
| original | BART | 4096 | 5.00E-05 | 0.9996 | 0.9716 |
| alternative | BART | 2048 | 5.00E-05 | **0.9997** | 0.7724 |
| alternative_2 | BART | 4096 | 1.70E-05 | 0.9994 | 0.6161 |
| alternative_3 | BART | 2048 | 1.70E-05 | 0.9995 | 0.6039 |
| original | vaswani_wmt_en_de_big | 4096 | 5.00E-05 | 0.9995 | 0.9508 |
| alternative | vaswani_wmt_en_de_big | 2048 | 5.00E-05 | **0.9997** | 0.6882 |
| alternative_2 | vaswani_wmt_en_de_big | 4096 | 1.70E-05 | 0.9992 | 0.1616 |
| alternative_3 | vaswani_wmt_en_de_big | 2048 | 1.70E-05 | 0.9991 | 0.1436 |

PD2
| *transformer name* | *architecture* | *max tokens* | *learning rate* | *SC-score* | *Coverage* |
| --- | --- | --- | --- | --- | --- |
| original | BART | 4096 | 5.00E-05 | 0.9922 | 0.2279 |
| alternative | BART | 2048 | 5.00E-05 | **0.9983** | 0.8167 |
| original | vaswani_wmt_en_de_big | 4096 | 5.00E-05 | 0.997 | 0.8352 |
| alternative | vaswani_wmt_en_de_big | 2048 | 5.00E-05 | **0.9978** | 0.6348 |

| PD3 |  |  |  |  |  |
| --- | --- | --- | --- | --- | --- |
| transformer name | architecture | max tokens | learning rate | SC-score | Coverage |
| original | BART | 4096 | 5.00E-05 | 0.7427 | 0.5976 |
| alternative | BART | 2048 | 5.00E-05 | **0.7644** | 0.5576 |
| original | vaswani_wmt_en_de_big | 4096 | 5.00E-05 | 0.8389 | 0.6464 |
| alternative | vaswani_wmt_en_de_big | 2048 | 5.00E-05 | **0.9362** | 0.511 |

**Table S2.** We trained both transformer architectures on the same amino-acid datasets and measured the accuracy and coverage.
| PD4 |  |  |  |  |  |
| --- | --- | --- | --- | --- | --- |
| transformer name | architecture | max tokens | learning rate | SP-score | Coverage |
| original | BART | 4096 | 5.00E-05 | 0.9785 | 0.5054 |
| alternative | BART | 2048 | 5.00E-05 | **0.9878** | 0.8405 |
| original | vaswani_wmt_en_de_big | 4096 | 5.00E-05 | 0.993 | 0.995 |
| alternative | vaswani_wmt_en_de_big | 2048 | 5.00E-05 | **0.9949** | 0.9794 |

**Table S3.**
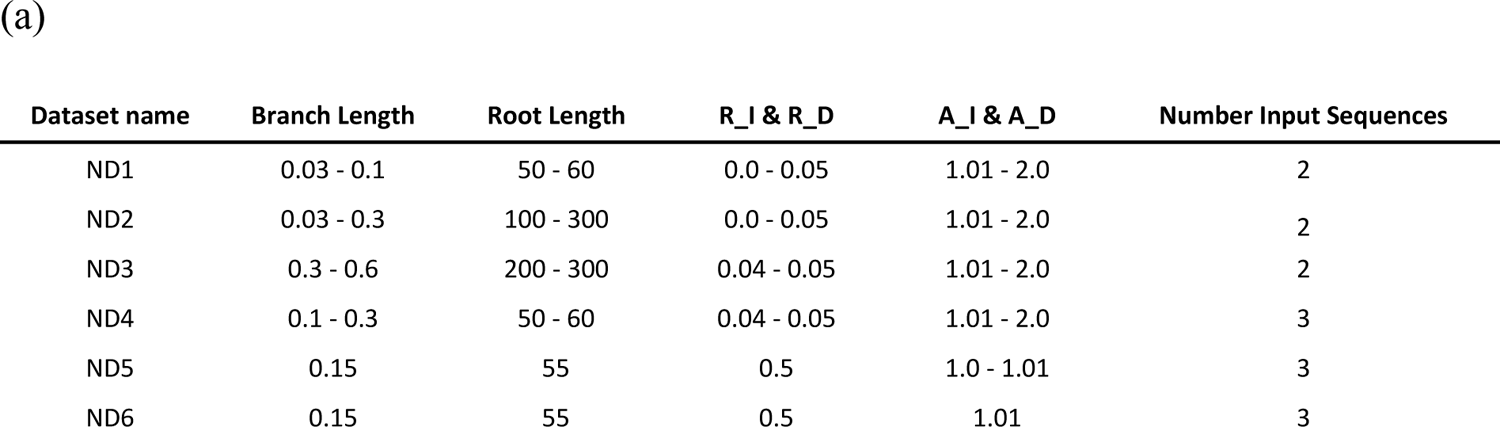
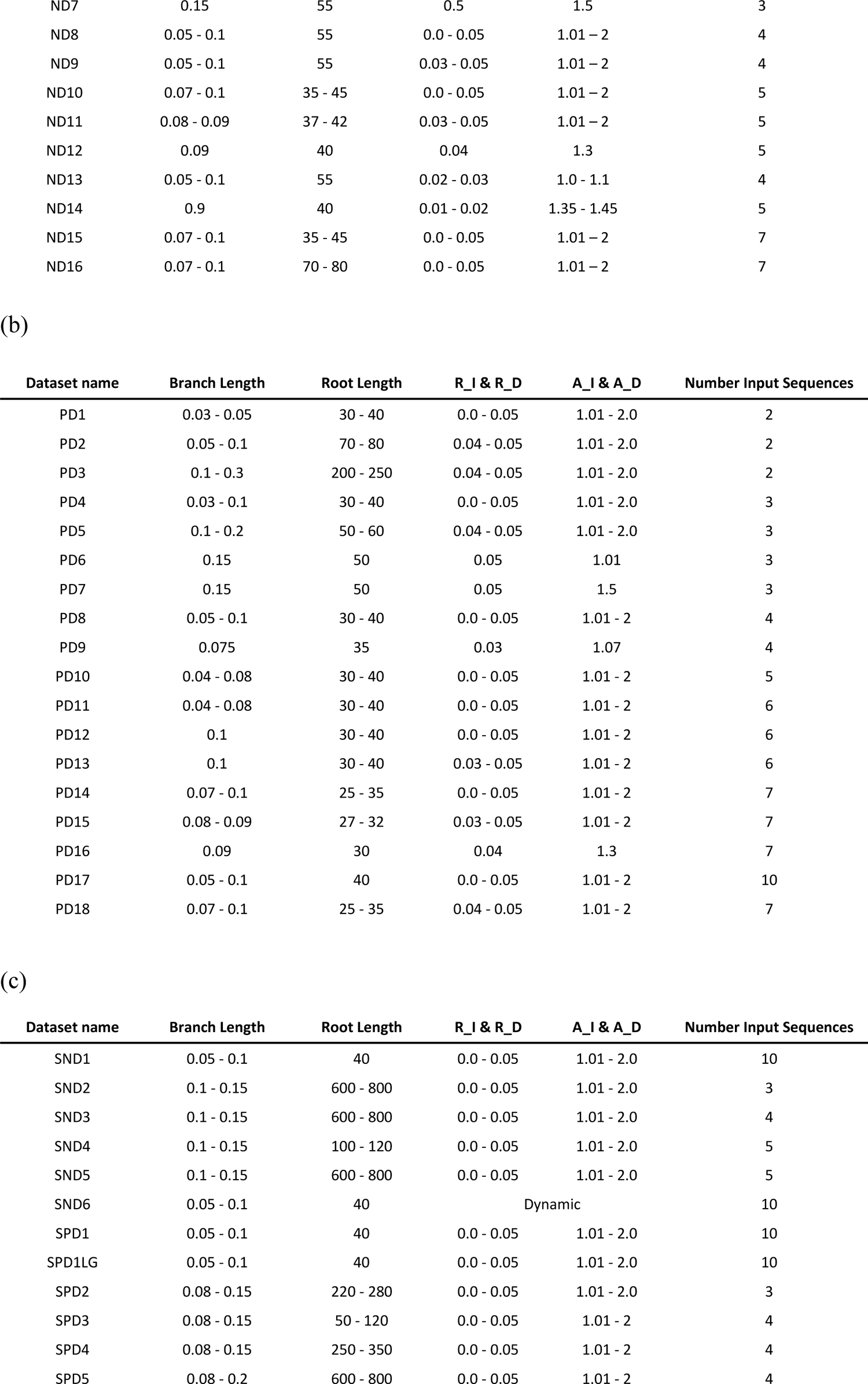
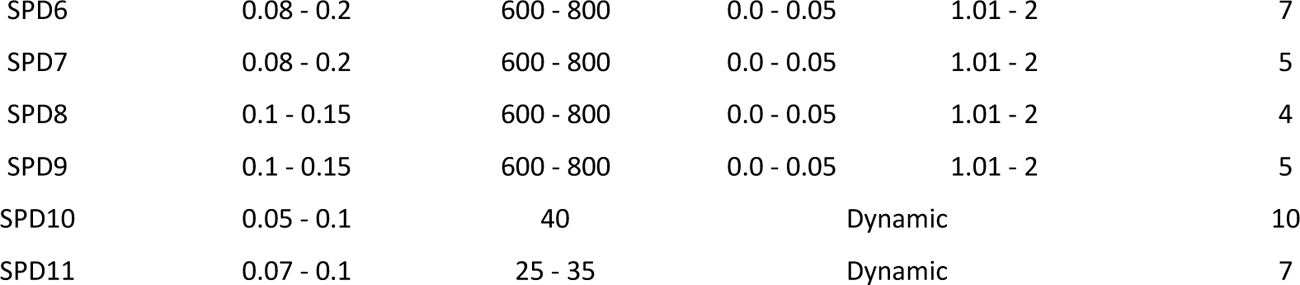
The table contains the specific information to generate the datasets used in this work. The R_I & R_D and the A_I & A_D refer to the indel parameters of the SpartaABC. The root length described in the table is the input of the SpartaABC, which the latter sample uniformly from the min range * 0.8 and the max range * 1.1. The datasets name refers to the order in which the transformers were trained on. For example, the first protein transformer was trained on dataset PD1, then the optimized weights were the starting point of PD2, etc. Similarly, the nucleotides transformer was first trained on ND1. Tables (a), (b) refer to nucleotide and protein datasets, respectively. Table (c) refers to special datasets (hence the “S” at the start of the dataset name). In this table, datasets SND1 and SPD1LG are the datasets used for creating *Fig. 2*. The other datasets, SND2 to SND5 and SPD2 to SPD9 refers to nucleotide and protein sequences used for the “segmented transformers”, respectively. SND6 and SPD10 refers to the misclassification datasets, and thus, the indel parameters are sampled differently for each “domain” (see “Generating Misspecification Dataset” section).

## Notes

### Competing Interest Statement

The authors have declared no competing interest.

